# Development and Characterisation of a Versatile Single-Domain Antibody Specific for M1-linked Ubiquitin Chains

**DOI:** 10.64898/2026.07.05.736589

**Authors:** Julian Koch, Shun-Je Bhark, Verian Bader, Berthe Katrine Fiil, Blanca López-Méndez, Josefine Bebe Rasthøj, Dominik Priesmann, Oscar Mejias-Gomez, Marta Braghetto, Guillermo Montoya, Mads Gyrd-Hansen, Konstanze F. Winklhofer, Steffen Goletz, Jonathan N. Pruneda, Rune Busk Damgaard

## Abstract

Ubiquitin signalling is mediated by structurally distinct polyubiquitin chains that encode discrete cellular functions. Progress in deciphering this ubiquitin code, particularly for the less abundant atypical chain types, has been hindered by limited availability of versatile chain type-specific affinity reagents. Here, we demonstrate that human single-domain antibodies (sdAbs) provide a versatile scaffold for the generation of ubiquitin linkage-specific binders. Using phage display and synthetic human sdAb libraries, we identified 2A6, an sdAb that specifically recognises methionine-1 (M1)-linked ubiquitin chains. To our knowledge, 2A6 represents the first reported sdAb with specificity for a defined homotypic ubiquitin chain linkage. 2A6 bound M1-linked ubiquitin chains with nanomolar affinity and was specific for M1-linked chains at the level of both diubiquitin and long polyubiquitin chains. AlphaFold3 modelling, supported by saturation mutagenesis, predicted that 2A6 recognises the proximal and distal ubiquitin moieties together with the region near the M1 linkage. Functionally, 2A6 enabled specific detection and enrichment of M1-linked ubiquitin across multiple applications, including ELISA, immunoblotting, immunoprecipitation under semi-denaturing conditions, substrate ubiquitination analysis, and immunofluorescence microscopy. The sdAb can be readily produced in *E. coli* from a single expression plasmid, providing a tractable, cost-effective and versatile reagent for investigating M1-linked ubiquitin signalling. Our work establishes sdAbs as a versatile scaffold for ubiquitin linkage-specific affinity reagents, providing a framework for the development of analogous binders specifically targeting additional ubiquitin linkages or architectures.

## Introduction

Posttranslational modification of proteins with ubiquitin (Ub) regulates virtually all aspects of cellular biology [1]. Ub is conjugated primarily to the ε-amino group in lysine (K) residues of the substrates on which polyUb chains are formed by additional ubiquitination of Ub itself on any of its seven internal lysines (K6, K11, K27, K29, K33, K48, or K63) or its N-terminal methionine (M1). These eight polyUb linkages form the basis of the “Ub code” [2]. The individual polyUb chains function as discrete signals in cells and can each control important cellular functions, ranging from proteasomal degradation to organelle quality control and non-proteolytic regulation of kinase signalling [1,2].

The basis of the functional diversity of the different polyUb linkages is that they are structurally distinct and adopt unique conformations. Chain types such as K48- and K6-linked Ub adopt compact and “closed” conformations, whereas K63- and M1-linked chains adopt more “open” and flexible conformations [3]. The spectrum of conformations each chain type can adopt is determined by the linkage residue. The position of the linkage point on the proximal Ub determines the relative orientation of the Ub being conjugated (the distal Ub) and hence the possible conformations and dynamics of that specific Ub linkage [3]. The different polyUb chains therefore display unique distributions of hydrophobic interaction patches, e.g. the isoleucine (I)44 patch, which can be bound by specific Ub-binding domain (UBD)-containing proteins that facilitate the specific functions for different polyUb chains.

Deciphering the “Ub code” is important for understanding the biology that is regulated by Ub-dependent signalling. K48- and K11-linked chains mainly target substrates for degradation in the 26S proteasome [4], and K63-linked chains mediate mostly non-degradative signalling downstream of immune receptors, in the DNA damage response, and in protein trafficking [5]. The remaining five linkage types are less abundant (together typically constituting <20% of the Ub linkages in cells), and cellular functions of these so-called “atypical” Ub linkages remain mostly elusive. K6-, K27-, K29-, and K33-linked chains have been linked to a variety of stress responses, including mitophagy, viral infection, and proteotoxic stress [1,6,7]. The functions of M1-linked chains are better understood. M1-linked chains (also known as “linear” Ub chains) are conjugated by the linear Ub chain assembly complex (LUBAC) [8–11] and cleaved by the deubiquitinase (DUB) OTULIN [12–14]. They are very lowly abundant in cells and only constitute ∼0.5% of cellular polyUb chains [15]. Yet, they have critical roles in regulation of innate immune signalling, particularly TNF receptor-1 [9–11,16] and NOD2 signalling [13,17], as well as emerging functions in stress responses and metabolism [18–21].

Beyond its established roles in protein homeostasis and signal transduction, Ub signalling is increasingly recognised as being regulated by additional layers of complexity. For example, lipids and small metabolites can modulate the stability and activity of E3s and DUBs, thereby influencing ubiquitination and signalling outputs [22,23]. Moreover, recent work has demonstrated that ubiquitination is not restricted to proteins, but can also target non-proteinaceous substrates, such as lipids, sugars, and ADP-ribose, further expanding the scope of Ub-dependent regulation and challenging traditional views of the Ub system [24–28]. These findings highlight the remarkable versatility of Ub signalling and underscore the need for robust and specific tools to dissect the mechanisms and functions of Ub signalling.

One of the main reasons for the limited insight into linkage type-specific functions of atypical polyUb chains is the scarcity of specific reagents for their efficient enrichment and detection. Specific or selective affinity tools have been reported for all Ub linkage types [29], including antibodies and antibody fragments (Fabs) [30–33], affimers [34], and engineered UBDs and DUBs [13,21,35,36]. However, only a limited range of reagents are available for atypical linkages (M1, K6, K27, K29, and K33), with M1-linked Ub being a relative exception [29]. For M1 linkages, available reagents include the 1E3 Fab/IgG [32], the M1-specific Ub binder (M1-SUB) based on the UBAN domain of NEMO [13], and the M1-Trap derived from catalytically inactive OTULIN [21]. Whereas 1E3 is primarily used for detection, M1-SUB and M1-Trap are not suitable for detection and are mainly used for enrichment. Generally, the availability, performance, and methodological compatibility of Ub linkage-specific affinity reagents vary considerably and rarely is a single reagent able to perform robustly across multiple applications. This complicates experimental design and may reduce confidence in the consistency and specificity of observations. Such limitations continue to constrain functional and mechanistic dissection of linkage-specific polyUb signalling, particularly of the atypical linkages.

Here, using M1-linked Ub as a model linkage, we demonstrate that single-domain antibodies (sdAbs) provide a viable scaffold for the generation of versatile, multi-purpose linkage-specific Ub affinity reagents. sdAbs are small antibody fragments of approximately 15-17 kDa derived from the variable domain of a heavy chain and consisting of a framework and three complementarity-determining regions (CDRs). Despite lacking a variable light chain, sdAbs can achieve affinities and specificities comparable to those of conventional Fab fragments [37–39]. Owing to their small size, sdAbs can access sterically restricted epitopes that are inaccessible or difficult for larger antibody formats, and they can readily be fused to functional modules, e.g. tags, making them versatile tools for detection, imaging, and targeted protein manipulation [37,40]. Using phage display and a set of synthetic human sdAb libraries, we identified and characterised 2A6, an sdAb that specifically recognises M1-linked Ub. To our knowledge, 2A6 represents the first reported sdAb with specificity for a defined homotypic Ub chain linkage type. 2A6 binds M1-linked diUb chains with high nanomolar affinity and is compatible with a broad range of applications, including enzyme-linked immunosorbent assay (ELISA), immunoblotting, immunoprecipitation under semi-denaturing conditions, and immunofluorescence microscopy. The sdAb can be readily produced in *E. coli* from a single expression plasmid, providing a tractable, cost-effective, and versatile addition to the experimental toolbox for investigating M1-linked Ub signalling. More broadly, our findings demonstrate that sdAbs can achieve Ub chain linkage specificity, establishing a framework for the development of analogous reagents targeting other Ub chain types or modifications.

## Results

### Selection of M1-specific sdAbs from a phage display library

We set out to select and develop M1 linkage-specific sdAbs to generate a tractable and versatile, multi-purpose reagent in a small, engineerable, and cost-efficient scaffold [37]. We devised a selection protocol using a set of synthetic, human sdAb libraries with a functional diversity of ∼10^10^ unique functional sdAb variants, designed based on position-specific CDR amino acid usage and varying CDR3 length (8 up to 23 amino acid residues) [39,41,42] (**Figure S1A**) for phage display to obtain sdAbs able to specifically bind M1-linked Ub (**Figure 1A**). To increase stringency and the specificity of the obtained binders, we included detergents (1% v/v IGEPAL CA-630, 0.5% v/v sodium deoxycholate (NaDC), and 0.1% v/v sodium dodecyl sulfate (SDS)) in the wash buffer after positive selection (**Figure 1A**). Through four rounds of negative and positive selection, we removed binders for monoUb, K48-linked polyUb, and K63-linked polyUb as the most abundant and likely off-targets and enriched for binders for M1-linked diUb (**Figure 1B**). Clonal screening of enriched sdAb-phages by ELISA revealed that a large proportion of binders demonstrated specificity or selectivity for M1-linked diUb over K48- and K63-linked diUb as well as monoUb (**Figure 1C**). Sequencing of all isolated sdAb clones revealed high variability among the clones in both CDR2 and CDR3 (CDR1 is invariant in the libraries), including no apparent similarity to the CDRs of the published 1E3 Fab [32] (**Figure S1B**). Interestingly, we only enriched clones with CDR3 lengths of 11 or 14 residues, and not any from the libraries with up to 23 residues in length (**Figure S1B**). Clones 2A6, 2C6, 2E4, and 2G6 were among the most promising binders, combining high ELISA signal with low reactivity towards monoUb and K48- or K63-linked chains (**Figure 1C**). We generated the variants 2A6.1 and 2G6.1 by swapping the CDR2s of 2A6 and 2G6 to test if it would result in improved binders (**Figure S1B**). We expressed selected FLAG-tagged sdAbs in *E. coli* and tested the specificity and sensitivity of the purified sdAbs towards M1-linked diUb. Immunoblotting using an HRP-coupled anti-FLAG secondary antibody indicated that the 2A6 clone was the most specific and sensitive binder (**Figure 1D**). 2A6.1 lost sensitivity compared with parental 2A6, and 2G6.1 started to show mild cross-reactivity towards K48- or K63-linked chains (asterisks), similar to 2C6 (**Figure 1D**). This was supported by direct ELISA using purified sdAbs (**Figure 1E**). Clone 2A6 detected 0.02 μg M1-linked diUb over 1 μg K48- or K63-linked diUb, demonstrating high specificity for M1 linkages and at least a 50-fold difference in apparent sensitivity between M1 and K63 or K48 linkages. In contrast, the more sensitive clone 2G6.1 did not discriminate between M1- and K63-linked diUb at the lowest M1-linked diUb concentration (0.02 μg) and therefore lacked comparable specificity (**Figure 1E**). In contrast, other clones identified in the monoclonal screen, including 2E4 and 2C6, displayed substantially lower sensitivity when tested as purified sdAbs by direct ELISA (**Figures 1C, E**). Interestingly, our clone screening also revealed that clone 2E11 was a good non-selective polyUb binder that did not discriminate between K48-, K63-, and M1-linked chains, but did generally not detect monoUb (**Figures 1C-E**). In conclusion, clone 2A6 was the most promising binder identified, and we continued to characterise it.

**Figure 1.**
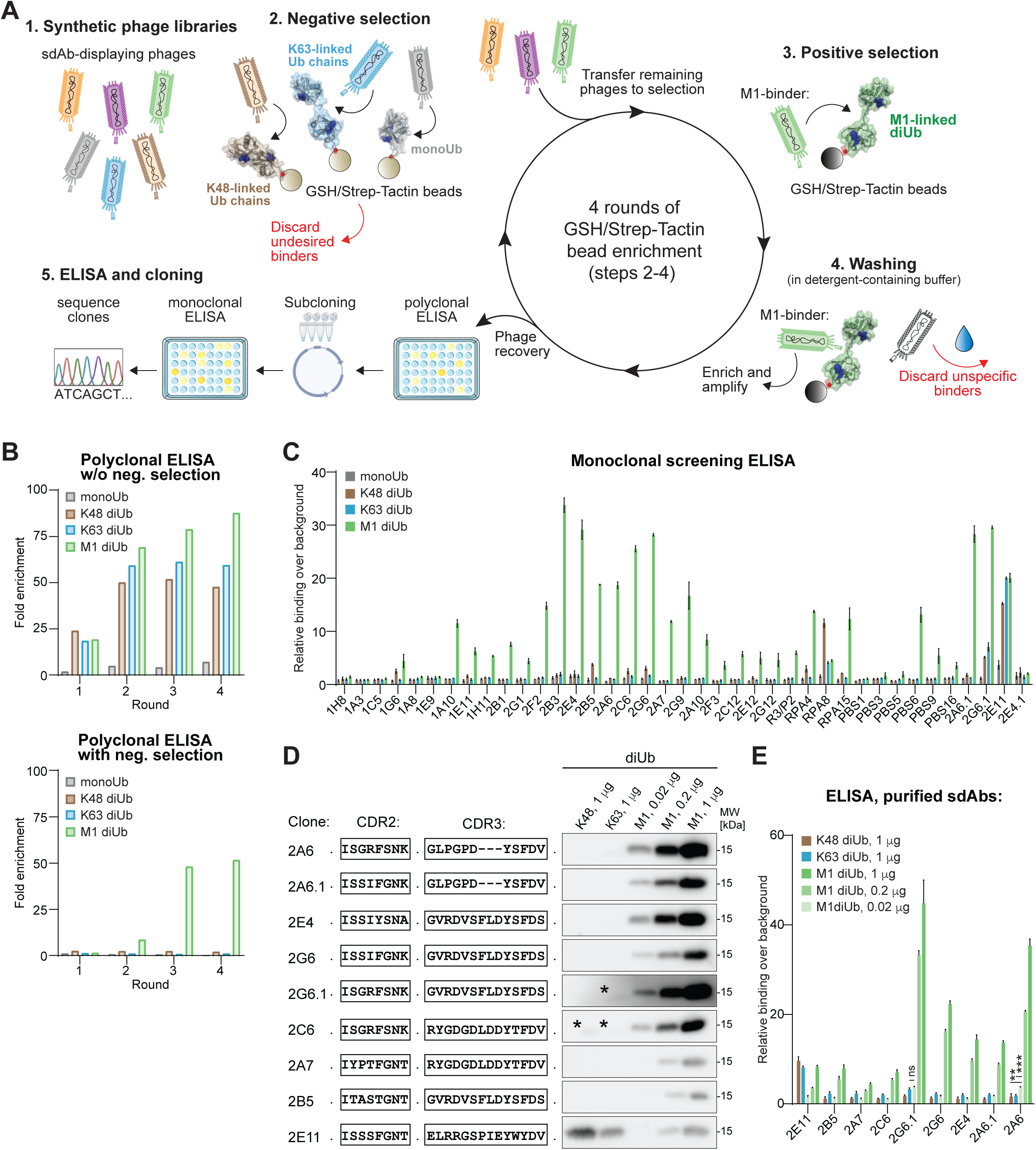
Phage display selection of sdAbs binding to M1-linked Ub. (A) Schematic overview of the sdAb selection protocol phage display using GST- and Strep-tagged Ub chains and GSH- and Strep-Tacting beads. (B) Phage ELISA showing the polyclonal enrichment of K48-, K63-, and M1-linked diUb-binding phage populations after each round of phage display selection, without (top) or with (bottom) negative selection. Plates were coated with 1 μg/well of mono or diUbs. (C) ELISA of selected sdAb clones expressed in the bacterial periplasm, showing binding to K48-, K63-, and M1-linked diUb. Plates were coated with 1 μg/well of mono or diUbs. Bars represent mean ± SD of three independent experiments. (D) CDR2 and CDR3 sequences of selected sdAb clones and corresponding immunoblots using FLAG-tagged sdAbs showing their binding to K48-, K63, and M1-linked diUb. Asterisks mark cross-reactivity of M1-selective sdAbs. (E) ELISA on selected, purified sdAbs showing their binding to K48-, K63, and M1-linked diUb. Bars represent mean ± SD of three independent experiments. Analysed by two-sided Student’s *t*-test. ns, no significant (p < 0.05), ** p < 0.01, *** p < 0.001.

### Biophysical characterisation and structural basis for 2A6 specificity

Our ELISA analyses indicated greater sensitivity of 2A6 for M1-linked Ub compared with K48- or K63-linked Ub. We therefore measured its binding affinity for M1-, K48-, and K63-linked chains. Isothermal titration calorimetry (ITC) revealed that 2A6 bound M1-linked diUb with a dissociation constant (*K_d_*) of 749 ± 117 nM and M1-linked tetraUb with a *K_d_* of 197 ± 37 nM (**Figure 2A, B**). Notably, we did not detect any apparent binding between 2A6 and K48-linked diUb or K63-linked diUb (**Figure 2C**). The ITC measurements also indicated the formation of a ∼1:1 sdAb:diUb complex and a ∼3:1 sdAb:tetraUb complex (N = 0.65 for M1-linked diUb and N = 0.22 for tetraUb), consistent with 2A6 binding across one M1 linkage. Further supporting this mode of binding, 2A6 was able to protect M1-linked Ub chains from hydrolysis by USP21 in solution in a concentration-dependent manner (**Figure S2A**).

**Figure 2.**
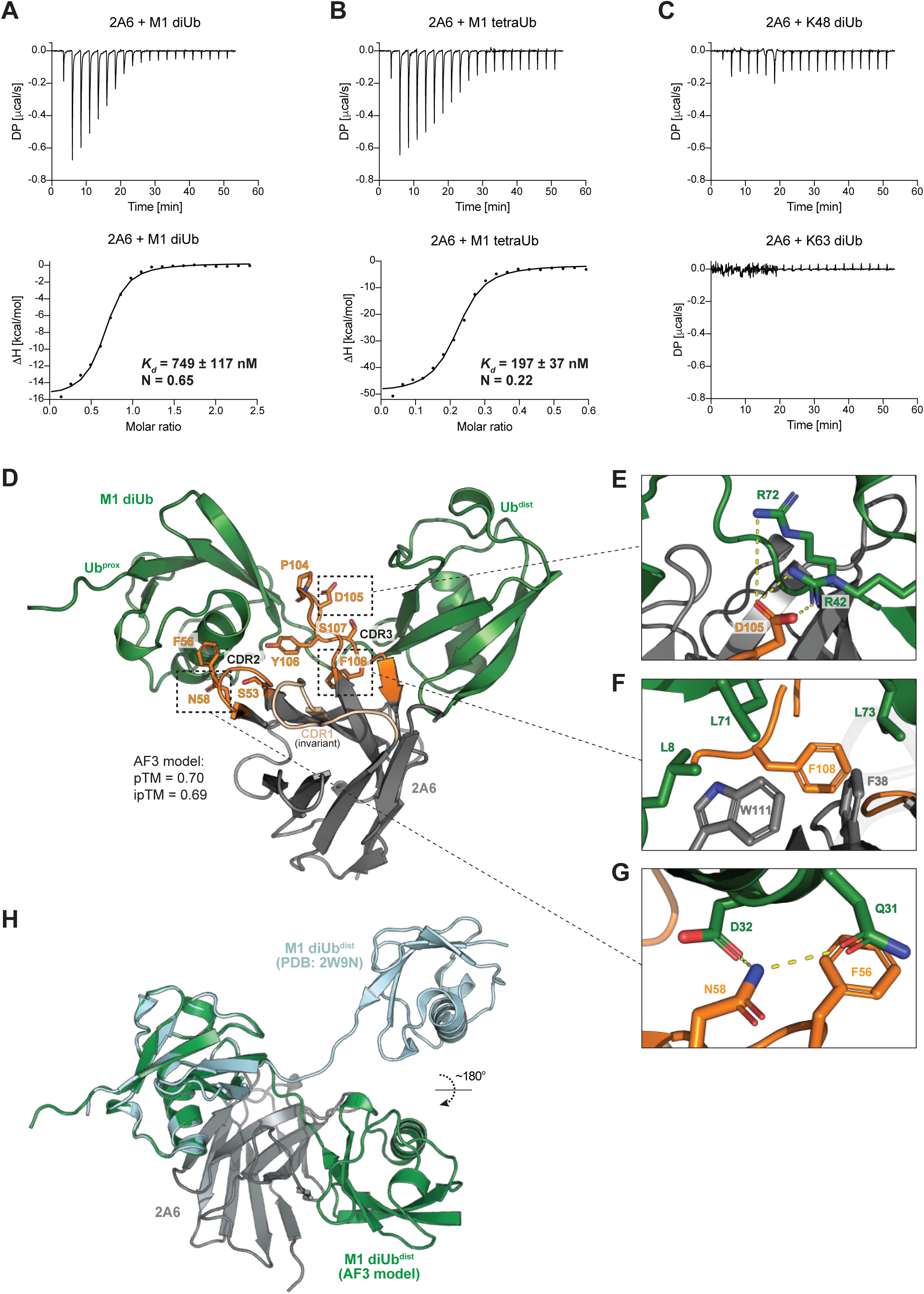
Biophysical and structural characterisation of M1-linkage specific sdAb 2A6. (A-B) ITC titration of sdAb 2A6 (28 μM, in cell) with (A) M1-linked diUb (350 μM in syringe) or (B) M1-linked tetraUb (80 μM in syringe), showing plots of raw heat (top) and derived isotherms (bottom) with fitted curve. *K_d_*’s are are shown ± the error of the fitting. (C) ITC titration of sdAb 2A6 (28 μM, in cell) with K48-linked diUb (350 μM in syringe) (top) or K63-linked diUb (350 μM in syringe), showing plots of raw heat. (D) AlphaFold3 (AF3) model of sdAb 2A6 (grey) binding to M1-linked diUb (green). CDR2 and CDR3 domains are highlighted in orange, with relevant amino acids shown as sticks. (E) Detailed view of the modelled interactions between 2A6 CDR3 residue D105 and distal Ub residues R42 and R72. The viewing orientation has been rotated slightly from Figure 2D to enhance visibility. (F) Detailed view of the modelled interactions between L8, L71, and L73 of the distal Ub with F108 of the 2A6 CDR3 region as well as W111 and F38 of the 2A6 backbone. The viewing orientation has been rotated slightly from Figure 2D to enhance visibility. Residues 99,101,106, and 110 from 2A6 have been hidden to enhance visibility. (G) Detailed view of the modelled interactions between D32 and Q31 of the distal Ub and N58 of the 2A6 CDR2 region. The viewing orientation has been rotated slightly from Figure 2D to enhance visibility. (H) Alignment of the proximal Ub (green) from the 2A6 AlphaFold3 model (AF3; grey) with the proximal Ub of 2W9N [44] (sky blue).

To elucidate the structural basis of binding and the M1 linkage specificity of 2A6, we attempted to crystallise the sdAb in complex with M1-linked diUb. However, 2A6 did not yield crystals, either in complex with M1-linked diUb or alone. We also attempted to crystallise two additional M1 linkage binders, 2C6 and 2G6, in complex with M1-linked diUb. Although neither formed crystals in complex with diUb, both crystallised in the absence of antigen (**Table S1** and **Figure S2B**). To obtain insights into the basis for specificity, we therefore modelled the interaction between 2A6 and M1-linked diUb using AlphaFold3 [43] (**Figures S2C-E**). The top-ranked model (**Figure 2D**) supported a plausible interface in which 2A6 binds between the proximal and distal moieties and across the M1 linkage, consistent with the stoichiometries indicated by our ITC measurements, with good-to-moderate confidence in overall complex assembly (pTM = 0.70) and inter-chain geometry (ipTM = 0.69). The independently generated models showed strong structural convergence, supporting a consistent binding mode despite some uncertainty in relative orientation (**Figures S2C-E**).

The top-ranked model predicted that D105 of 2A6’s CDR3 forms a salt bridge with R42 and R72 of the distal Ub (**Figure 2E**). In addition, extensive hydrophobic interactions between F38, W48, F108, and W111 of 2A6 and L8, L71, and L73 of the distal Ub near the M1 linkage suggested favourable burial of these residues at the binding interface (**Figure 2F**). Contacts with the proximal Ub were also observed, with N58 of CDR2 engaging Q31 and D32 (**Figure 2G**). Together, these interactions suggest that 2A6 recognises both the distal and proximal Ub moieties, as well as the region near the M1 linkage.

Intriguingly, the predicted conformation of M1-linked diUb in complex with 2A6 differed markedly from that of free M1-linked diUb [44] (**Figure 2H**). Superimposition on the proximal Ub revealed that binding to 2A6 was predicted to be associated with an ∼180° rotation and 26 Å displacement of the distal Ub, resulting in a pronounced bend in the M1-linked chain (**Figure 2H**). This rearrangement enables the relatively small 2A6 sdAb to engage both the distal and proximal Ub moieties as well as area near the M1 linkage to achieve specificity.

To improve the affinity of 2A6 for M1-linked Ub, we performed saturation mutagenesis at selected positions. However, analysis of the permitted variants (**Figure S2F**) revealed that none of them enhanced binding to M1-linked diUb compared with parental 2A6 (**Figure S2G**). Notably, several residues predicted by the AlphaFold3 model to contribute to binding either did not tolerate substitution or permitted only conservative substitutions. These included W48, D105 and F108, as well as neighbouring residues, and to a lesser extent F38, N58, and W111 (**Figure S2F**). Together, these findings support the AlphaFold3-predicted interface between 2A6 and M1-linked Ub and suggest that the parental 2A6 already adopts a near-optimal binding configuration that is not readily improved by single amino acid substitutions.

### *In vitro* validation of 2A6 for detection and enrichment of M1-linked Ub

We then sought to validate 2A6 for the detection and enrichment of M1-linked Ub. Immunoblotting using FLAG-tagged 2A6 demonstrated that the sdAb retained its specificity for M1-linked Ub when tested against the full panel of diUb linkages (**Figure 3A**). 2A6 detected 0.02 μg M1-linked diUb but failed to detect 1 μg of any other linkage type, indicating at least a 50-fold preference for M1-linked chains. In contrast, the control sdAb 2E11 displayed little linkage selectivity and detected all diUb linkages, although the signal was weaker for K27-linked diUb and slightly stronger for K48-linked diUb (**Figure 3A**). This specificity was maintained on a panel of longer polyUb chains, with 2A6 exclusively recognising M1-linked chains, whereas 2E11 detected all chain types tested (**Figure 3B**).

**Figure 3.**
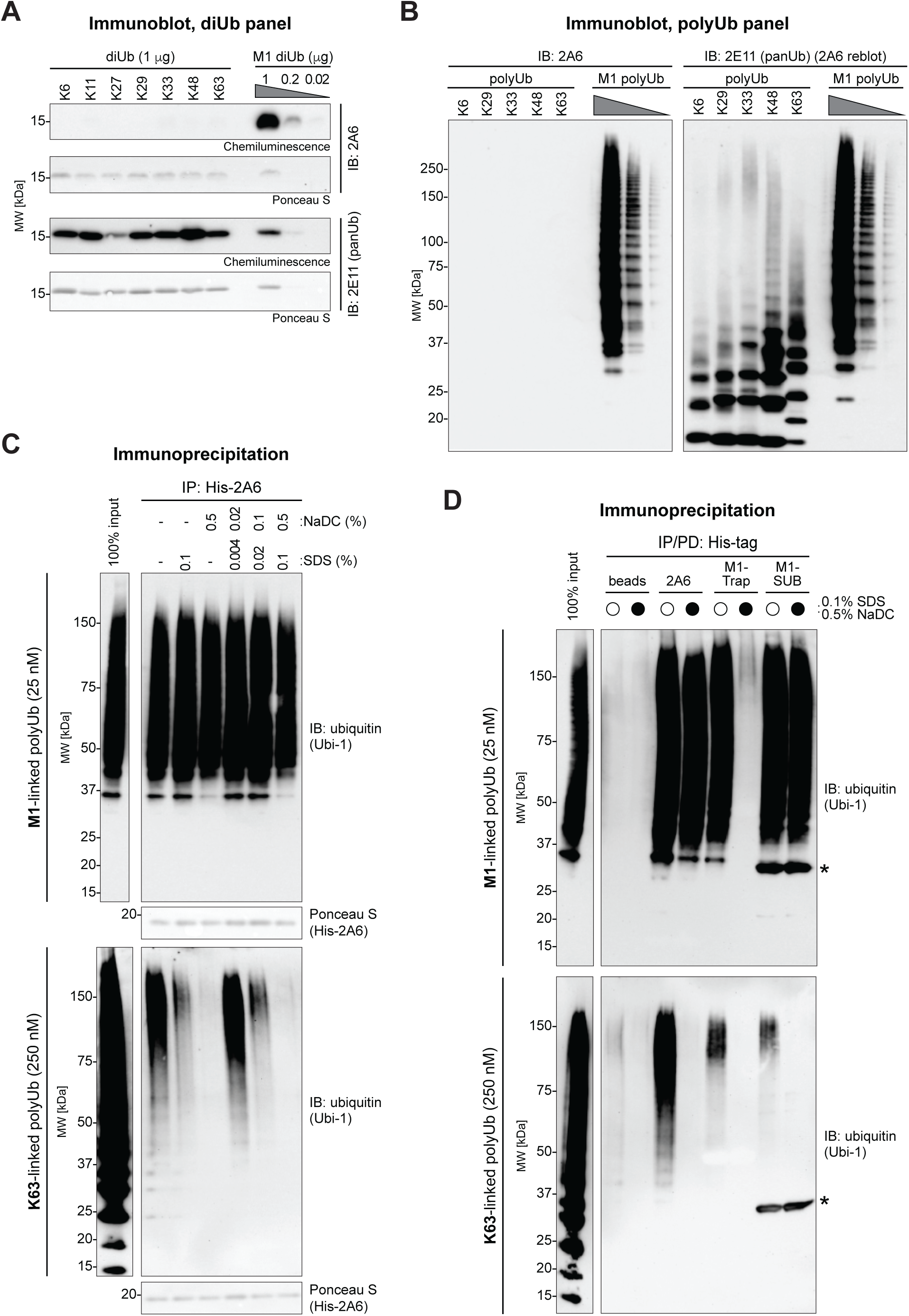
Validation and applications of sdAb 2A6 specificity for M1-linked Ub *in vitro*. (A) Immunoblots using FLAG-tagged sdAbs 2A6 and 2E11 on a diUb panel. Ponceau S-stained membranes show diUb input. (B) Immunoblots using FLAG-tagged sdAbs 2A6 and 2E11 on a polyUb panel. 2E11 was used to reblot on membranes first incubated with 2A6. (C) Immunoblots of immunoprecipitation (IP) of recombinant polyUb chains by His-tagged sdAb 2A6 in the absence or presence of NaDC and SDS detergents. Ponceau S-stained membranes show sdAb IP levels. (D) Immunoblots of immuno- or affinity purification of polyUb chains using His-tagged sdAb 2A6, M1-Trap, and M1-SUB in the absence or presence of NaDC and SDS detergents as indicated. IP levels of sdAb 2A6, M1-Trap, and M1-SUB are in Figure S3A.

Next, we investigated whether 2A6 could be used to immunoprecipitate M1-linked chains. To assess its selectivity, we compared recovery of M1- and K63-linked polyUb, using K63-linked chains at a tenfold higher molar concentration to mimic the substantially greater abundance of K63- than M1-linked chains in cells [15]. His-tagged 2A6 efficiently recovered M1-linked chains ranging from tetraUb to species exceeding 150 kDa (>20 Ub moieties) (**Figure 3C**). Under our standard immunoprecipitation conditions (50 mM Tris-HCl pH 7.4, 1% IGEPAL CA-630, 150 mM NaCl, and 5 mM MgCl₂), 2A6 also recovered K63-linked chains, although primarily species larger than 75 kDa (>10 Ub moieties), suggesting non-specific recognition of higher-order structures within long K63-linked chains. Consistent with this, 2A6 recovered virtually all M1-linked chains present in the reaction but only a fraction of the K63-linked chains (**Figure 3C**; compare IP lanes and 100% input lanes), consistent with the substantially lower affinity of 2A6 for K63-linked chains observed in ITC. Because the phage display selections were performed using wash buffers containing the detergents NaDC and SDS, and because the M1-specific 1E3 Fab/IgG requires high concentrations of urea to achieve linkage-selective immunoprecipitation [32], we tested whether detergent supplementation could improve the selectivity of 2A6. Addition of either NaDC or SDS markedly reduced recovery of K63-linked chains, with NaDC having the greater effect under the conditions tested, while their combined use (0.5% NaDC and 0.1% SDS) abolished K63-linked chain recovery (**Figure 3C**). In contrast, the recovery of M1-linked chains was largely unaffected, demonstrating that 2A6 can specifically immunoprecipitate M1-linked chains under appropriately stringent, semi-denaturing conditions.

We next benchmarked 2A6 against the established M1-linked Ub enrichment reagents M1-SUB and M1-Trap [13,21]. Under these *in vitro* immunoprecipitation conditions, all three reagents recovered M1-linked chains as well as varying amounts of K63-linked chains, with 2A6 exhibiting the highest level of K63-chain recovery (**Figures 3D and S3A**). Supplementation of the buffer with NaDC and SDS abolished recovery of K63-linked chains by both 2A6 and M1-SUB, while preserving recovery of M1-linked chains, indicating that both reagents can achieve specific enrichment of M1-linked chains under stringent detergent conditions. In contrast, the M1-Trap did not tolerate these detergent conditions, resulting in a complete loss of binding to both M1- and K63-linked chains (**Figures 3D and S3A**).

Together, these results demonstrate that 2A6 is a highly selective reagent for the detection and enrichment of M1-linked Ub. In addition, the non-selective sdAb 2E11 may provide a useful pan-polyUb reagent.

### Cellular applications of 2A6 for investigating M1-linked Ub signalling

Having established the specificity of 2A6 *in vitro*, we next tested its performance in cellular assays, beginning with immunoprecipitation of M1-linked polyUb from cells. Because wild-type (WT) cells contain very low levels of M1-linked chains at steady state [15,45], we compared WT and OTULIN knockout (KO) cells, which accumulate high levels of M1-linked Ub [46]. Consistent with this, 2A6 efficiently enriched M1-linked chains from lysates of mouse embryonic fibroblasts (MEFs), with markedly greater recovery from OTULIN-KO than WT cells (**Figure 4A**), demonstrating that 2A6 can selectively enrich M1-linked Ub from complex biological samples. Similar results were obtained in human HCT116 cells, where 2A6 efficiently enriched M1-linked chains from OTULIN-deficient cells (**Figure 4B**). In contrast, the non-selective sdAb 2E11 enriched polyUb independently of OTULIN status, further supporting the selectivity of 2A6 (**Figure S4A**). Importantly, enrichment by 2A6 was completely abolished in OTULIN-HOIP double-KO cells, confirming that recovery was dependent on LUBAC activity and therefore represented M1-linked chains (**Figure 4B**). This selectivity was observed both in the standard immunoprecipitation buffer and in buffer supplemented with NaDC and SDS detergents. Notably, the detergent-containing buffer reduced background recovery, consistent with the increased stringency observed in immunoprecipitations of purified Ub chains under the same conditions (**Figure 4B**).

**Figure 4.**
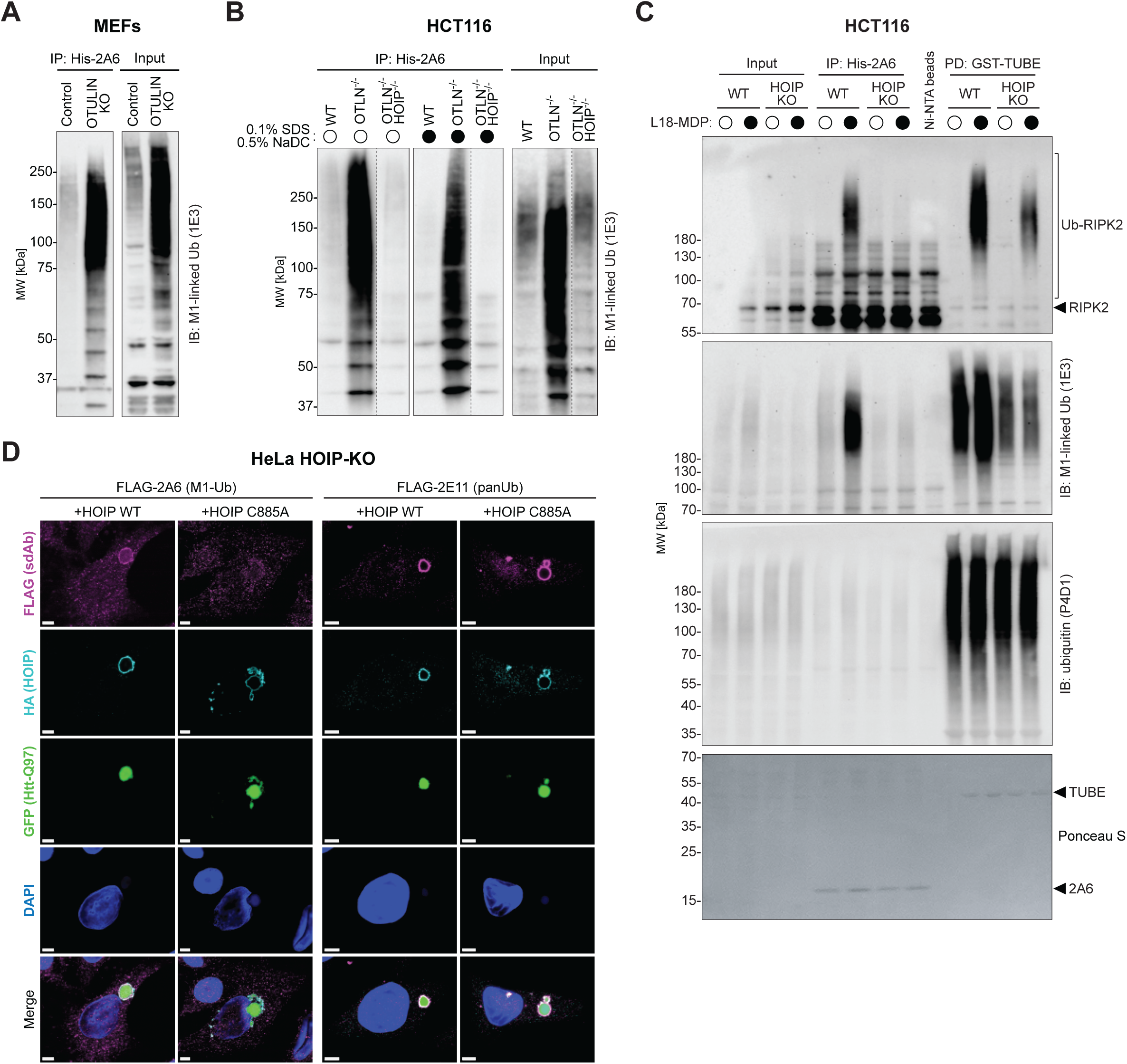
Applications of 2A6 for investigating M1-linked Ub signalling in cells. (A) Immunoblots of IP with His-tagged sdAb 2A6 from control (OTULIN^+/-^) and OTULIN-KO MEFs in IP buffer without NaDC and SDS detergents. (B) Immunoblots of IP with His-tagged sdAb 2A6 from WT, OTULIN^-/-^, or OTULIN^-/-^-HOIP^-/-^HCT116 cells in the absence or presence of NaDC and SDS detergents. (C) Immunoblots of IP with His-tagged sdAb 2A6 or pulldown with GST-TUBE from WT or HOIP-KO HCT116 cells treated with L18-MDP as indicated. IP and pulldowns were performed in the presence of NaDC and SDS detergents. Ponceau S-stained membranes show sdAb IP and TUBE pulldown levels. (D) Immunofluorescence microscopy analysis of sdAbs 2A6 and 2E11 co-localising with Htt-Q97 aggregates. HeLa HOIP-KO cells co-expressing Htt-Q97-GFP (green) and either HOIP^WT^ or HOIP^C885A^ (cyan) were stained with FLAG-tagged sdAbs (magenta). DNA was stained with DAPI (blue). Scale bars, 5 μm.

Next, we investigated whether 2A6 could be used to analyse substrate ubiquitination in addition to global cellular levels of M1-linked Ub. In the NOD2 signalling pathway, LUBAC ubiquitinates the adaptor protein RIPK2 following receptor stimulation [13,17,47]. WT and HOIP-KO cells were stimulated with the NOD2 ligand L18-MDP, after which M1-linked Ub was enriched using 2A6 and total Ub using non-selective tandem Ub-binding entities (TUBEs) [47,48] under stringent, detergent-containing conditions. Immunoblotting for RIPK2 following enrichment with 2A6 revealed robust and entirely HOIP-dependent ubiquitination of RIPK2 in response to L18-MDP stimulation (**Figure 4C**), showing that 2A6 specifically enriches RIPK2 species ubiquitinated with M1-linked chains. A similar pattern was observed following TUBE enrichment; however, HOIP deletion did not completely abolish RIPK2 ubiquitination, consistent with RIPK2 being modified by other Ub chain types in addition to M1-linked chains, including K63 linkages [47,49], which are recognised by the TUBE but not by 2A6. In addition, 2A6 readily distinguished WT and HOIP-KO cells at the level of M1-linked chain enrichment. Although M1-linked chains were detectable in unstimulated WT cells, their abundance increased markedly following L18-MDP stimulation. In contrast, enrichment of M1-linked chains was strongly reduced in HOIP-KO cells. Notably, the signal recovered by 2A6 from unstimulated WT cells exceeded that observed in unstimulated HOIP-KO cells (**Figure 4C**, compare lanes 5 and 7), demonstrating that 2A6 is sufficiently sensitive to detect and enrich M1-linked Ub even at very low steady-state levels.

We next investigated whether 2A6 could be used to monitor M1-linked Ub signalling *in situ*. Aggregation of misfolded proteins, including mutant Huntingtin (Htt) with an elongated poly-glutamine stretch (Htt-polyQ) associated with Huntington’s disease, promotes recruitment of LUBAC and subsequent M1-linked ubiquitination of the aggregates [50]. We therefore examined whether 2A6 could detect M1-linked Ub accumulation at Htt-97Q aggregates by confocal fluorescence microscopy. In HeLa HOIP-KO cells reconstituted with HOIP^WT^, 2A6 signal accumulated at Htt-Q97 aggregates together with HOIP (**Figure 4D**). In contrast, reconstitution with the catalytically inactive HOIP^C885A^ mutant resulted in no clear accumulation of 2A6 signal at the aggregates, despite HOIP^C885A^ localising to Htt-Q97 aggregates and robust accumulation of signal from the non-selective sdAb 2E11 under the same conditions (**Figure 4D**). These findings indicate that multiple Ub chain types are conjugated to Htt aggregates, but that accumulation of the 2A6 signal is specifically dependent on LUBAC-catalysed M1-linked ubiquitination.

Collectively, these data demonstrate that 2A6 enables specific detection, enrichment, and analysis of M1-linked ubiquitination in biochemical, cellular, and imaging-based applications.

## Discussion

In this study, we identified and characterised 2A6, a versatile, multi-purpose human sdAb that specifically recognises M1-linked Ub chains with high nanomolar affinity. To our knowledge, 2A6 is the first reported sdAb with specificity for a defined homotypic Ub chain linkage, establishing sdAbs as a viable scaffold for the generation of linkage-selective Ub affinity reagents. Interestingly, a recent study by Yogesh Kulathu and colleagues described NbSL3.3Q, a camelid sdAb (nanobody) identified through yeast surface display, that specifically recognises K48/K63-branched Ub chains [51]. Our work extends these findings by showing that sdAbs can also discriminate between the eight homotypic Ub chain linkages. While NbSL3.3Q recognises a complex K48/K63-branched triUb species by contacting all three Ub moieties, achieving picomolar affinity, 2A6 recognises only two Ub moieties yet discriminates M1-linkages from the seven remaining homotypic linkage types. Together, these studies demonstrate that sdAbs can achieve remarkable specificity towards both complex Ub architectures and individual homotypic Ub chain linkages, highlighting their considerable potential as a versatile scaffold for the development of next-generation Ub affinity reagents. Furthermore, our findings highlight the functional diversity of the human sdAb libraries, demonstrating that highly specific Ub linkage-selective sdAbs can be isolated directly without the need for affinity maturation.

To gain insight into the structural basis of M1-linkage recognition by 2A6, we combined AlphaFold3 modelling with saturation mutagenesis. Together, these approaches support a model in which 2A6 engages both the proximal and distal Ub moieties and simultaneously contacts the area near the M1 linkage itself. This binding mode differs substantially from that of endogenous, M1-linkage-specific UBAN domains of ABIN proteins and NEMO, which function as molecular rulers that recognise the distance between I44 of the distal Ub and F4 of the proximal Ub, subtly bending the chain compared with the conformation of free M1-linked diUb, to preferentially recognize M1-linked chains [52,53]. In contrast, our model predicts that binding of 2A6 is accompanied by a pronounced reorientation of the distal Ub, allowing the compact sdAb scaffold to bridge the two Ub moieties and directly bind the linkage region. Interestingly, the predicted interaction of 2A6 with M1-linked diUb resembles that of the M1-specific 1E3 Fab [32] in that both binders engage the proximal and distal Ub moieties while also contacting the region surrounding the M1 linkage. This apparent convergence suggests that simultaneous recognition of both Ub moieties and the linkage itself may represent a general principle underlying highly selective recognition of M1-linked Ub chains, despite the fundamentally different architectures of Fab and sdAb scaffolds. Importantly, residues in 2A6 predicted to make key contacts with M1-linked diUb were generally highly intolerant to substitution in saturation mutagenesis, providing independent support for the proposed interface and suggesting that the parental 2A6 already adopts a near-optimal binding configuration.

Existing M1-linked Ub-specific affinity reagents have transformed the field but differ considerably in their optimal applications [29]. Whereas the 1E3 Fab/antibody [32], and commercialised variants of it, is primarily used for detection in immunoblotting and immunofluorescence microscopy and M1-SUB and M1-Trap have been developed principally as enrichment reagents [13,21], 2A6 proved compatible with a broad range of experimental approaches, including ELISA, immunoblotting, immunoprecipitation, substrate enrichment, and immunofluorescence microscopy. Notably, 2A6 maintained specificity under semi-denaturing immunoprecipitation conditions containing SDS and NaDC, enabling stringent enrichment of M1-linked chains while reducing non-specific interactions. This may be particularly valuable for proteomics applications, where increased stringency is expected to minimise co-purification of unmodified interaction partners and thereby improve discrimination between directly ubiquitinated substrates and associated proteins [29,54]. In addition to 2A6, the non-selective sdAb 2E11 may also represent a useful reagent for pan-polyUb detection and enrichment. Together with straightforward bacterial production from a single expression plasmid, these properties make 2A6 a practical and versatile addition to the experimental toolbox for investigating M1-linked Ub signalling.

Beyond providing a new reagent for studying M1-linked ubiquitination, our work establishes a conceptual framework for developing sdAbs against additional Ub chain architectures, including homotypic chains. The small size, stability, and engineerability of sdAbs make them particularly well suited for applications that are challenging for conventional antibodies, including intracellular expression as intrabodies, fusion to fluorescent or enzymatic reporters for live-cell imaging, targeted intracellular protein manipulation, and proximity-labelling approaches. Engineered UBDs and DUBs, such as the UBAN domain of NEMO and catalytically inactive OTULIN, can also be adapted for intracellular sensing and manipulation of M1-linked Ub [13,21,35]. However, because these reagents are derived from endogenous signalling proteins, they retain interaction surfaces and regulatory features that may perturb native protein interactions and signalling pathways. In contrast, sdAbs represent orthogonal affinity reagents with few, if any, endogenous binding partners beyond their intended target. Consequently, they are less likely to interfere with cellular signalling through off-target interactions, making them particularly attractive scaffolds for intracellular sensing and modulation of ubiquitin signalling. This concept is not restricted to M1-linked Ub but is likely to extend to sdAbs targeting other Ub linkages or architectures, enabling selective intracellular sensing and manipulation of the Ub code while minimising the risk of unintended perturbation of endogenous signalling networks. sdAbs therefore represent an attractive scaffold for the development of specific reagents for the less well-characterised atypical Ub chains and emerging modifications such as ester-linked chains [55], helping to overcome one of the major technical barriers to deciphering the Ub code and the biology it regulates.

## Materials and Methods

Key materials and reagents used in this study are listed in **Table 1**.

**Table 1:**
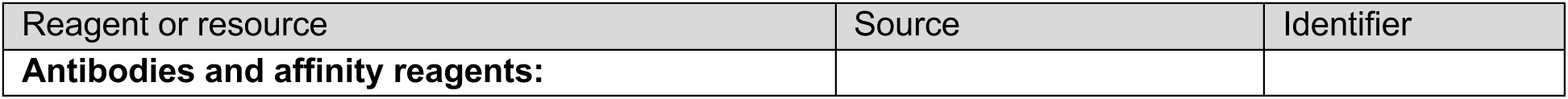
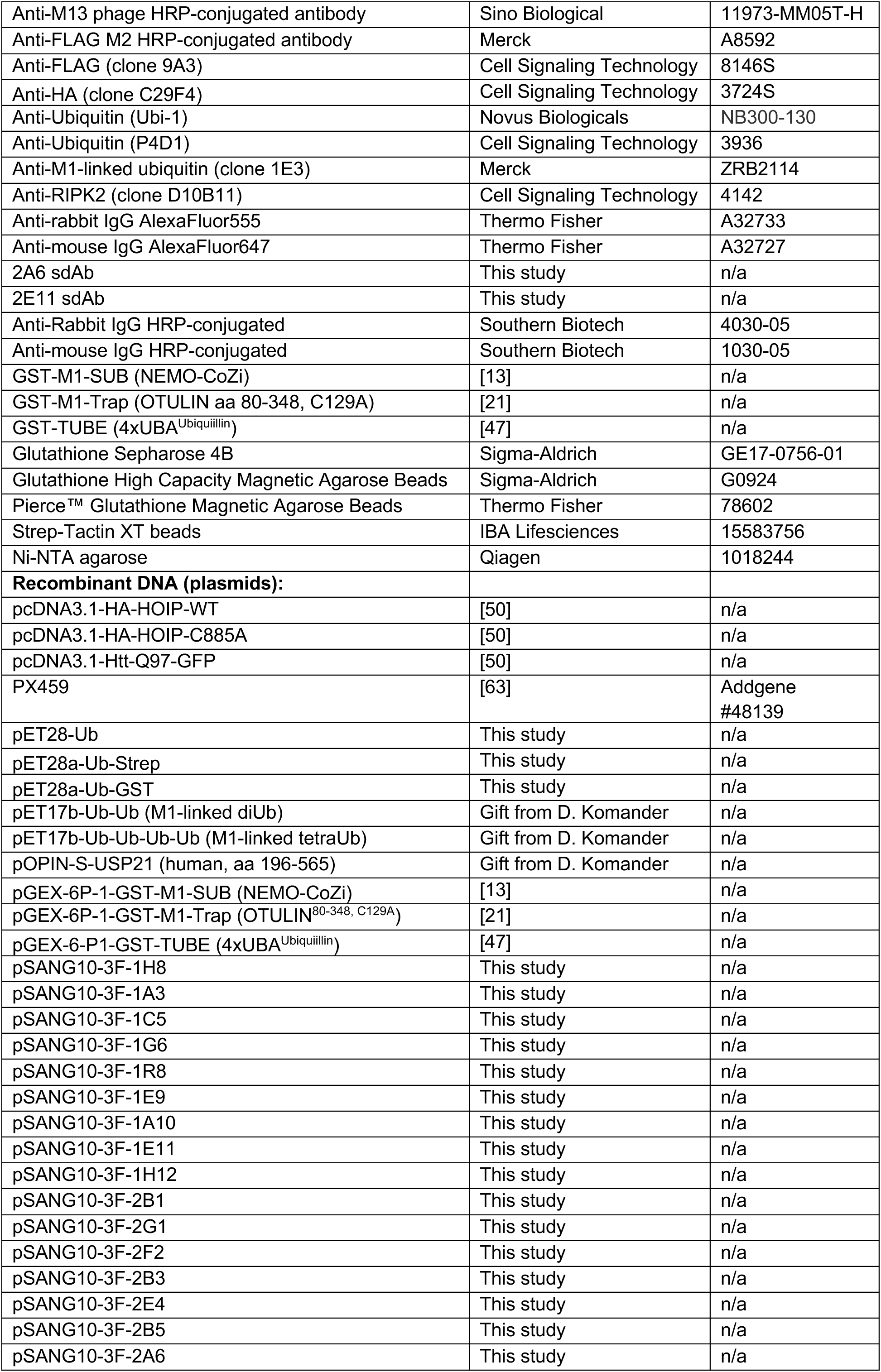
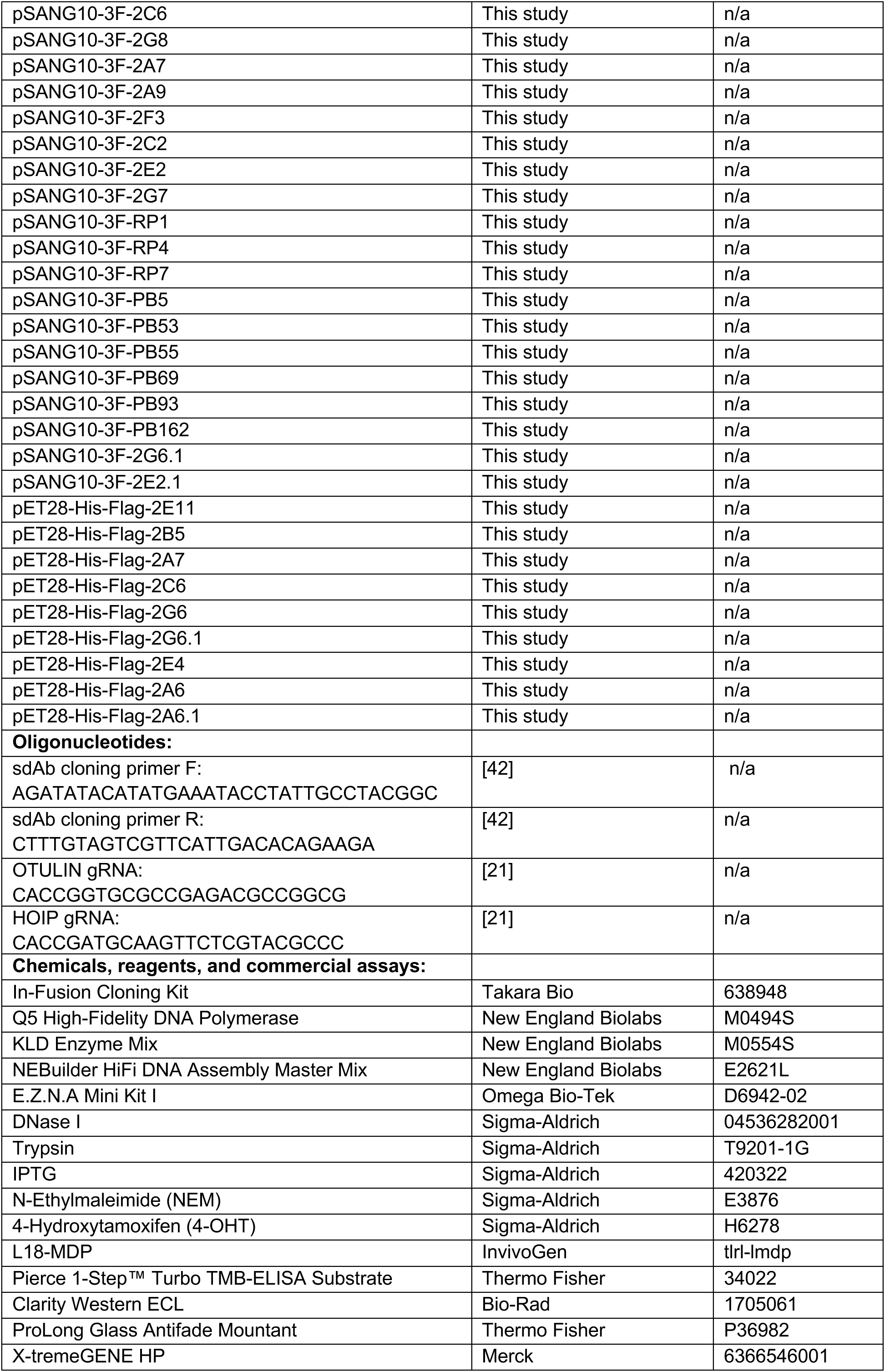
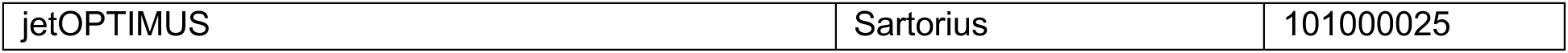
List of key reagents and resources, including their identifiers, used in this study.

### Plasmids and cloning

Desired inserts were either PCR-amplified from available constructs or synthesized and obtained commercially (Twist Bioscience). Purified PCR products were cloned into linearised pSANG10-3F, pET28a, pcDNA3.1, or pGEX-6P-1 vectors using In-Fusion primers and cloning kit (Takara Bio). Site-directed mutagenesis via overlap extension PCR was done using the Q5® High-Fidelity DNA Polymerase (New England Biolabs; NEB) and the NEBaseChanger for primer design. For both cloning and mutagenesis, transformation was performed by heat-shocking into Stellar™ Competent *E. coli* for 60 seconds at 42°C, followed by plating on LB agar containing kanamycin (50 μg/mL) or ampicillin (100 μg/mL). Single colonies were picked and expanded in LB media with the respective antibiotic, and plasmids were purified using the E.Z.N.A. Mini Kit I (Omega Bio-tek). Regions of interest were sequenced by Sanger sequencing (Macrogen Europe) using primers targeting the T7 or CMV promoters and a custom pGEX reverse sequencing primer (5’-CCGGGAGCTGCATGTGTCAGAGG-3’). Plasmids used in this study are listed in **Table 1**.

### Phage display

Phage display selections were performed using a mix of four sdAb M13 phage libraries based on the human HEL4 scaffold [56], which had been further engineered for increased stability and low aggregation propensity, as well as variation in CDR3 length (8-23 amino acid residues) [39,41,42]. The libraries were generated using the trypsin-sensitive helper phage KM13 that allows a gross-reduction of background via trypsin-based elution and inactivation of background phages [57]. Selections were carried out as previously described [42], with minor modifications. TG1 *E. coli* (Agilent) were streaked in advance on a M9 minimal media plate without antibiotics. For negative selection (NS), GST- or Strep-tagged monoUb and K48- and K63-linked polyUb (>tetraUb) was immobilized on magnetic glutathione beads (Sigma-Aldrich) or Strep-Tactin™XT beads (IBA Lifesciences) in 500 μL of PBS (10 mM P_i_ pH 7.4, 138 mM NaCl, 2.7 mM KCl) for 1 h at room temperature (RT), tumbling overhead. The tags used for negative selection were alternated between panning rounds, using Strep/Strep-Tactin™XT in round 1 and 3 and GST/glutathione in round 2 and 4. Amounts of tagged monoUb and polyUb loaded into each immobilization reaction was incrementally increased over four rounds of negative selection: monoUb, 1-4 nmol; K48-linked polyUb, 0.6-6 nmol; K63-linked polyUb, 3-30 nmol. For positive selection (PS), GST-or Strep-tagged M1-linked diUb was immobilized on beads as described above. The immobilisation tags were alternated in the opposite pattern to the negative selection, using GST/glutathione in rounds 1 and 3 and Strep/Strep-Tactin™XT in rounds 2 and 4. The amount of tagged M1-linked diUb loaded into the immobilization reaction was incrementally decreased through four rounds of positive selection, from 2-0.5 nmol. After washing both the loaded NS- and PS-beads with PBS, they were incubated with a mixture of ∼1×10^12^ M13 phages from the mixed libraries [39,41,42] in 500 μL of 2% (w/v) milk powder in PBS (mPBS) for 1 hour at RT, tumbling overhead. Subsequently, the NS-beads were precipitated and removed. The mPBS containing the depleted libraries was transferred onto the loaded PS-beads, incubating 1 h at RT with tumbling overhead. The PS-beads were washed ten times with 1 mL of RIPA buffer (50 mM Tris-HCl pH 7.4, 1% IGEPAL CA-630, 150 mM NaCl, 5 mM MgCl2, 1 mM EDTA, 0.1% (w/v) SDS, 0.5% (w/v) deoxycholate) and then two times with 1 mL of PBS, or ten times with PBS with 0.1% Tween-20 (PBS-T) and then two times with PBS, depending on the screening campaign. Phages were afterwards eluted with 500 μL of 1 mg/mL trypsin (Sigma-Aldrich) for 10 min at RT, tumbling overhead. The trypsinised phages were used to infect 5 mL of exponentially growing TG1 *E. coli* (OD600 of approx. 0.4) in 2xYT containing 1% w/v glucose and 100 μg/mL carbenicillin for 30 min. The cultures were then plated on 2xYT agar plates containing 1% w/v glucose and 100 μg/mL carbenicillin overnight. The number of colonies was counted, and the plates were scraped and used to inoculate 10 mL 2xYT with 100 μg/mL carbenicillin. After growing to OD600 of 0.4, 2.5×10^10^ M13 helper phages were added and left to incubate for 30 min before pelleting the infected TG1 *E. coli* by centrifugation, then resuspending in 2xYT with 100 μg/mL carbenicillin and 50 μg/mL kanamycin, growing overnight. The overnight culture was cleared by centrifugation at 5000 x*g* and the supernatant was mixed with PEG/NaCl (20% PEG6000, 2.5M NaCl) at a dilution factor of 4:1 (supernatant:PEG/NaCl) to precipitate phages and centrifuged at 10,000 x*g*. The phage pellet was washed and number of phage particles per mL was quantified photometrically as described in [58]. Phages were cryoprotected with 2xYT with 20% (v/v) glycerol, frozen, and stocked for subsequent rounds of selection or analysis. Enrichment and specificity of binders for each round was confirmed in a polyclonal ELISA against diUb (K48-, K63- and M1-linked), monoUb, and background-binding to milk using the phage stocks from each round and detecting with an anti-M13 phage HRP-coupled antibody (Sino Biological).

### Cloning of sdAbs from phagemids into expression vectors

After panning round 4, TG1 *E. coli* were infected with enriched phages as described above, then streaked on 2xYT plates containing 1% w/v glucose and 100 μg/mL carbenicillin. After growth, the plates were scraped and the phagemid was purified from the pellet using the E.Z.N.A. Mini Kit I (Omega Bio-tek, D6942). The region coding for the sdAbs was then amplified by PCR with flanking primers, cloned into the pSANG10-3F vector using the NEBuilder HiFi DNA Assembly Master Mix (New England Biolabs), transformed into NEB10 *E. coli* (New England Biolabs), and streaked on 2xYT plates containing 1% w/v glucose, 100 μg/mL streptomycin and 50 μg/mL kanamycin. Single clones were picked and sequenced.

### Screening and selection of monoclonal sdAbs

After transformation into BL21(DE3) *E. coli* (Agilent) with pSANG10-3F-sdAb-encoding plasmids (for secretion into the periplasm) and growth on a 2xYT agar plate with 50 μg/mL kanamycin, single colonies were picked and inoculated in 600 μL of 2xYT with μg/mL kanamycin in a 96-deepwell plates to make starter cultures. The starter cultures were used to inoculate a fresh 96-deepwell plate containing 600 μL of 2xYT with 50 μg/mL kanamycin. Expression was induced in the deep well plate after reaching OD600 of approx. 0.4 with 0.1 mM IPTG. After expression for 16 h at 25 °C, plates were centrifuged and the sdAb-containing supernatant from each well was then transferred to a immunosorbent plate for ELISA (see below) against M1-linked diUb using the anti-FLAG M2 antibody (Merck) for detection. sdAb expression was confirmed by dot blot, using the same detection antibody. Binders with high ELISA signal and visible expression on dot blot were re-expressed and tested for specificity in a second round of ELISA against M1-linked, K48-linked, K63-linked diUb, and monoUb. Binders of interest were re-grown for plasmid purification to identify sdAb sequences via Sanger sequencing.

### Enzyme-Linked Immunosorbent Assay (ELISA)

For ELISA, MaxiSorp Immuno Plates (Thermo Scientific) were coated with 0.01-1 μg M1-linked diUb, 1-2 μg of monoUb, K48-linked diUb, or K63-linked diUb per well in a volume 100 μL in sterile PBS overnight at 4 °C. After adsorbing Ub in the wells, they were blocked with 300 μL of 2% mPBS for 1 h at RT. For polyclonal ELISA on phages, 10 μL of the phage precipitate was mixed with 90 μL of 2% mPBS and added to the wells. For ELISA directly on sdAb-containing bacterial lysates, the blocked wells were re-filled with 50 μL mPBS before 50 μL of the cleared lysate from sdAb expressions were added. For ELISA on purified sdAbs, the sdAbs were added in 2% mPBS at a concentration of 1 μg/mL in a volume of 100 μL. All reactions were incubated for 1 h at room temperature. Plates were then washed three times with PBS-T (PBS, 0.1% v/v Tween-20) and incubated for 1 h at room temperature with either anti-M13 phage (Sino Biological) or anti-FLAG M2 (Merck) HRP-coupled antibodies before washing again three times in PBS-T and developing with 100 μL Pierce 1-Step™ Turbo TMB-ELISA Substrate (Thermo Fisher). Reactions were stopped by adding 100 μL 1 M HCl and the colorimetric signal was quantified by 450/650 nm absorbance on a Victor Nivo microplate reader (Revvity).

### Purification of sdAbs, Ub affinity reagents, and USP21

His-FLAG-tagged sdAbs were expressed from pET28a in SHuffle T7 Competent *E. coli* (New England Biolabs). The SHuffle strain is a strict requirement when expressing without the pelB leader sequence. M1-SUB (GST-NEMO-CoZi-His) [13], M1-Trap (GST-OTULIN^80-348;C129A^-His) [21], and TUBE (GST-4xUBA^Ubiquiillin^-His) [47] were expressed from pGEX-6P-1 in BL21(DE3) *E. coli*. His-SUMO-USP21 was expressed from pOPIN-S in BL21 CodonPlus(DE3) RIPL *E. coli* (Agilent Technologies). Chemically transformed bacteria were grown to an OD600 of 0.6 in LB broth supplemented with antibiotics, induced with 0.1 mM IPTG or 0.25 mM for USP21, and expressed overnight at 25 °C or 18 °C for USP21. Bacteria were harvested, pelleted, and flash frozen. For immobilized metal affinity chromatography (IMAC), pellets were resuspended in Buffer A (25 mM Tris pH 8.0, 200 mM NaCl, 20 mM Imidazole) supplemented with protease inhibitor cocktail (Sigma-Aldrich P8465) and 10 U/ml rDNase I (Sigma-Aldrich) for sonication. After sonication, pelleting (29,000 g, 20 min, 4 °C), and filtration, the cleared lysates were applied to a HisTrap FF Column (Cytiva). Protein was eluted by gradient up to 100% IMAC Buffer B (25 mM Tris pH 8.0, 200 mM NaCl, 500 mM Imidazole). Proteins were further purified by size exclusion chromatography (SEC) on a Superdex 75 Increase 10/300 GL SEC column (Cytiva), which consistently reduced non-specific cross-reactivity of the purified sdAbs in downstream assays. For USP21, pooled USP21-containing fractions were exchanged into PBS supplemented with 1 mM DTT and incubated overnight at 4°C with SENP1 protease to remove the SUMO-tag, followed by purification by size-exclusion chromatography. After buffer-exchanging into storage buffer (e.g. PBS), measuring UV-absorption at 280 nm, and adjusting protein concentration by MWCO units, purified proteins were aliquoted and stored at -80 °C.

### Purification of GST-tagged Ub

Ub-GST and GST-tagged M1-linked polyUb were expressed from pGEX-6P-1 or pET17b in BL21(DE3) *E. coli* by IPTG induction (0.1 mM IPTG) at an OD600 of 0.6 and expressed overnight at 25 °C. Bacteria were harvested, pelleted, and flash frozen. Pellets were then resuspended in PBS supplemented with protease inhibitor cocktail (Sigma-Aldrich P8465) and 10 U/ml rDNase I (Roche) for sonication. After sonication, pelleting (29,000 g, 20 min, 4 °C), and filtration, cleared lysates were incubated with 0.5 mL pre-washed slurry of Glutathione Sepharose 4B beads (Cytiva, 17075601) at RT for 1 h while tumbling slowly in a 49 mL Econo-Column (BioRad 7374251). After releasing the flowthrough, beads were washed three times with ice cold PBS and eluted via addition of five times 1 mL GSH elution buffer (PBS, 50 mM GSH, 1 mM DTT), incubating 1 min at RT each. Proteins were further purified on a Superdex 75 Increase 10/300 GL SEC column (Cytiva). After buffer-exchanging into storage buffer (e.g. PBS), measuring UV-absorption at 280 nm, and adjusting protein concentration by MWCO units, purified proteins were aliquoted and stored at -80 °C.

### Purification of Ub via perchloric acid precipitation and ion exchange (IEX)

MonoUb, M1-linked di-, tri, and tetraUb, and Ub-Strep constructs were expressed from pET28a or pET17b in BL21(DE3) *E. coli* by autoinduction using modified ZYM-5052 medium [59] for 24 h at 37 °C. Bacteria were harvested, pelleted and flash frozen. Pellets were resuspended in Bacterial Lysis Buffer (25 mM Tris pH 7.6, 150 mM NaCl, 1 mM EDTA) supplemented with protease inhibitor cocktail (Sigma-Aldrich P8465) and 10 U/ml rDNase I (Roche) for sonication. After sonication, pelleting (29,000 g, 20 min, 4 °C), and filtration, cleared lysates were acidified by dropwise addition of 70% perchloric acid to a final concentration of 0.5%. The mixture was stirred on ice for 30 min, cleared by centrifugation and dialysed against IEX Buffer A (50 mM sodium acetate pH 4.5) overnight at 4°C. The dialysed protein was loaded on a HiTrap SP FF (Cytiva) and eluted with by gradient up to 100% IEX Buffer B (50 mM sodium acetate pH 4.5, 500 mM NaCl). Proteins were further purified on a Superdex 75 Increase 10/300 GL SEC column (Cytiva). After buffer-exchanging into storage buffer (e.g. PBS), measuring UV-absorption at 280 nm, and adjusting protein concentration by MWCO units, purified proteins were aliquoted and stored at -80 °C.

### Purification of untagged sdAbs via Protein A

Untagged sdAbs for crystallisation were expressed from pSANG10-3F in BL21(DE3) pLysS *E. coli* (Thermo Fisher) by IPTG induction (0.1 mM IPTG) at an OD600 of 0.6 and expressed overnight at 25 °C. After expression, bacteria were lysed by sonication directly in expression medium supplemented with protease inhibitor cocktail and 10 U/ml rDNAse I. After pelleting (29,000 g, 20 min, 4°C) and filtration, the medium/lysate was diluted 1:1 with 2x binding buffer (25 mM NaH_2_PO_4_ and 15 mM Na_2_HPO_4_, pH 7.0) and loaded on a HiTrap Protein A HP Column (Cytiva). The sdAbs were eluted with a 0.1 M sodium citrate and instantly neutralized with 1 M Tris, pH 8.5. After buffer-exchanging into PBS and adjusting concentration by MWCO, proteins were aliquoted and stored at -80 °C.

### Assembly and purification of tagged and untagged Ub chains

All diUb and polyUb chains were assembled and purified as previously described [60]. Strep-and GST-tagged K48- and K63-linked Ub chains were assembled enzymatically using the same protocol [60], starting with Strep- or GST-tagged monoUb (C-terminally taggged) and then with stepwise addition of untagged monoUb over the course of the assembly to favour formation of chains containing a tag. Initially, one part of untagged monoUb was added to three parts of C-terminally tagged monoUb. Over the course of 6 h of enzymatic assembly, one part untagged monoUb was added every 30 min, adding up to twelve parts of untagged monoUb in total to the initial three parts of tagged monoUb. All assembled Ub chains were purified on a Superdex 75 Increase 10/300 GL SEC column (Cytiva) and separated into di-, tri-, tetra-, and >tetra Ub chains.

### SDS-PAGE and immunoblotting

For SDS-PAGE and immunoblotting, samples were resolved on NuPAGE Novex Bis-Tris 4-12% gels (Thermo Fisher) or self-cast 15% Bis-Tris gels. Gels were either stained in Coomassie Blue (0.1% Coomassie R-250, 40% ethanol, 10% acetic acid) and destained in water or proteins were transferred to 0.2 μm nitrocellulose membranes for immunoblotting using the Trans-Blot Turbo system. Membranes were stained with Ponceau S to verify transfer efficiency and equal loading, then blocked in 5% milk in PBS-T. After washing with PBS-T, membranes were incubated overnight at 4°C with primary antibodies (1:1000 dilution in 3% BSA, 0.01% NaN₃ in PBS-T) or sdAbs (1 μg/mL solution in 2% mPBS-T). The following day, membranes were washed three times with PBS-T and incubated with HRP-conjugated secondary antibodies for 1 h at RT. For immunoblots using sdAbs, FLAG-tagged variants (including 2A6 and 2E11) were used as primary antibodies together with an HRP-coupled anti-FLAG (M2 clone, Merck) as the secondary antibody. After three additional washes, blots were developed using Clarity Western ECL substrate (Bio-Rad) and imaged on a Bio-Rad ChemiDoc Imager. The Image Lab software (Bio-Rad) was used for image assembly and analysis. Antibodies are listed in **Table 1**.

### DUB protection assay

Deubiquitinase assay using USP21 on M1-linked tetraUb was performed as previously described [61]. Increasing concentrations of sdAb 2A6 (1.5-15 μM) were added to the reactions. USP21 activity was stopped at respective timepoints by the addition of 4x Laemmli sample buffer (LSB) to reach a 1x concentration (50 mM Tris pH 6.8, 10% glycerol, 100 mM DTT, 2% SDS, 0.1% Bromophenol blue), and the reactions were analysed by SDS-PAGE and Coomassie staining.

### Cell culture

Mouse embryonic fibroblasts (MEFs), HeLa cells, and HCT116 cells were maintained at 37°C in a humidified atmosphere containing 5% CO_2_. Heterozygous ERT2-*Cre*-*Otulin*^-/flox^ MEFs were described previously [15] and were cultured in phenol red-free Dulbecco’s Modified Eagle’s Medium (DMEM) (Gibco, 31053044) supplemented with 10% foetal bovine serum (FBS) (Biowest, S181B), 1% penicillin/streptomycin, GlutaMAX™, and pyruvate. OTULIN-KO MEFs were generated by culturing ERT2-*Cre*-*Otulin*^-/flox^ MEFs in the presence of 100 nM 4-hydroxytamoxifen (4-OHT; Sigma-Aldrich) for 7 days after which they were cultured in complete culture medium. Vehicle-treated (ethanol) heterozygous ERT2-*Cre*-*Otulin*^-/flox^ MEFs were used as controls. HeLa cells were cultured in DMEM with high glucose, GlutaMAX™, and pyruvate (Gibco, 31966047). HCT116 cells were cultured in McCoy’s 5A medium (Gibco, 36600088) or DMEM (Gibco, 31053044) supplemented with 10% FBS (Biowest, S181B) and 1% penicillin/streptomycin. HeLa and HCT116 HOIP-KO cells were described previously [47,62]. All cells were regularly tested for mycoplasma contamination.

### Generation of knockout cells using CRISPR-Cas9

For the generation of HCT116 OTULIN knockout (KO) and OTULIN-HOIP double-knockout cells, HCT116 WT or OTULIN-KO cells were grown to ∼60% confluency and then transfected with gRNAs targeting human OTULIN (5’-CACCGGTGCGCCGAGACGCCGGCG-3’) or human HOIP (5’-CACCGATGCAAGTTCTCGTACGCCC-3’) cloned into the pSpCas9(BB)-2A-Puro (PX459) backbone using X-tremeGENE HP (Merck) at a 2:1 (v/w) ratio according to the manufacturer’s instructions. 24 hours post-transfection, puromycin selection was initiated (1 μg/mL). Following five days of selection, surviving cells were allowed to recover in DMEM supplemented with 10% FBS (Biowest, S181B) and 1% penicillin/streptomycin without puromycin for 24 hours before they were trypsinised and plated into 96-well plates and grown from single clones in medium with 20% FBS and 1% penicillin/streptomycin. After ∼2 weeks of culture, clones were screened by immunoblotting. Individual clones were maintained in puromycin-free complete medium. PX459 was a kind gift from Professor Feng Zhang (Addgene plasmid #48139) [63].

### Immunoprecipitation and pulldown of purified Ub chains

For immunoprecipitation and pulldown, His-tagged 2A6 at a final concentration of 300 nM were combined with polyUb at a final concentration of 25 nM (M1-linked) or 250 nM (K63-linked). The proteins were added to 10 μl (slurry) pre-washed Ni-NTA agarose beads (Qiagen) per condition in a reaction volume of 1 mL per condition. Buffers used were base buffer (50 mM Tris-HCl pH 7.4, 1% IGEPAL CA-630, 150 mM NaCl, and 5 mM MgCl2) or RIPA-like buffer (base buffer + 0.1% (w/v) SDS, 0.5% (w/v) deoxycholate). After tumbling overhead for 1 h at RT, beads were washed three times with 800 μL of either RIPA-like buffer or base buffer and a pipette tip with a manually constricted orifice was used to fully aspirate washing buffer between washes. Bound material was eluted in a mixture of 4x LSB and IMAC buffer B before analysis by SDS-PAGE and immunoblotting.

### Immunoprecipitation of Ub and ubiquitinated proteins from cell lysates

MEFs and HCT116 cells cultured as described above were washed once with ice-cold PBS, then harvested and lysed on ice using base buffer or RIPA-like buffer supplemented with 5 mM N-ethylmaleimide (NEM) and cOmplete™ protease inhibitor cocktail (Roche). Lysates were cleared by centrifugation at 13,000 rpm for 10 minutes at 4°C, and supernatants were transferred to clean tubes and supplemented with 1 mM DTT. Lysates were then added to 10 μL (slurry) pre-washed Ni-NTA agarose beads (QIAGEN) before adding 0.5-2 μg of the respective sdAb, filling up to 500 μL with buffer if needed, and incubating for 16 h tumbling overhead at 4°C. Beads were washed three times with 800 μL of either ice-cold base buffer or RIPA-like buffer, respectively. A pipette tip with a manually constricted orifice was used to fully aspirate buffer between washes. Bound material was eluted in a mixture of 4x LSB and IMAC buffer B.

### Analysis of RIPK2 ubiquitination

Ub conjugates were enriched from cell lysates prepared from HCT116 WT and HOIP-KO cells [47]. Cells were seeded in 10 cm² dishes and treated with 200 ng/mL L18-MDP (Invivogen, tlrl-lmdp) for 1 hour as described previously [17]. After treatment, cells were lysed in RIPA-like lysis buffer (50 mM Tris-HCl, pH 7.4; 1% IGEPAL CA-630 (Merck); 150 mM NaCl; 5 mM MgCl₂; 0.5% deoxycholate; 0.1% SDS) supplemented with 20 mM imidazole, 1 mM DTT, 5 mM N-ethylmaleimide (NEM), and cOmplete™ protease inhibitor cocktail (Roche). Cleared lysates (10 min centrifugation at 21,000 × g, 4°C) were split in two: one for enrichment of M1-linked Ub with His-2A6 and one for enrichment of total Ub conjugates using GST-TUBE. For 2A6 enrichment, lysates were incubated overnight at 4°C with pre-washed Ni-NTA Agarose (Qiagen) and 2 μg 2A6 sdAb per sample on a rotating wheel. For the beads-only control, lysates from all four conditions were pooled equally and incubated with beads without 2A6. After enrichment, beads were washed six times in lysis buffer. Bound material was eluted using IMAC buffer B (Tris-HCl, pH 8; 200 mM NaCl; 500 mM imidazole) and Laemmli SDS sample buffer (Thermo Fisher). For total Ub conjugate enrichment, 15 μg of GST-TUBE was added per sample to magnetic glutathione beads (Thermo Fisher) and pre-incubated for 1-3 hours at 4°C. Beads were washed and lysates were added to incubate overnight at 4°C on a rotating wheel. Beads were then washed three times in lysis buffer, and bound material was eluted in 2× Laemmli SDS sample buffer (Thermo Fisher). Samples were resolved on NuPAGE Novex 4-20% Tris-Glycine gels (Thermo Fisher), transferred to 0.45 μm PVDF membranes using the Trans-Blot Turbo system, and analysed by immunoblotting. Membranes were stained with Ponceau S (Sigma-Aldrich) to assess 2A6 and GST-TUBE levels, transfer efficiency, and equal loading.

### Saturation mutagenesis

For saturation mutagenesis, 22 positions in 2A6 were selected based on the AlphaFold3 model. Mutants were generated from pSANG10-3F-2A6 by PCR using Q5 High-Fidelity DNA Polymerase (New England Biolabs) and primers containing NNK degenerate codons at the selected positions (Wells 1985). Primers were synthesised by TAG Copenhagen. Following PCR amplification, template DNA was removed and the products circularised using the KLD Enzyme Mix (New England Biolabs) prior to transformation into Stellar competent *E. coli*. For each NNK mutagenesis library made for each of the 22 positions, 96 colonies were selected, and the resulting variants were expressed in *E. coli* RIPL cells (Invitrogen) and screened by ELISA. Variants producing signals ≥90% of that obtained with parental 2A6 were subjected to Sanger sequencing.

### Isothermal titration calorimetry (ITC)

sdAbs and diUb or tetraUb chains were buffer-exchanged into degassed ITC buffer (10 mM HEPES pH 7.3, 150 mM NaCl). Protein concentrations were adjusted to 28 μM for the sdAbs, 320-380 μM for diUb, and 80 μM for tetraUb. ITC measurements were performed at 25 °C using a MicroCal Auto-iTC200 instrument (Malvern Panalytical). Ub chains at ∼350 μM concentration were loaded into the syringe and titrated into the calorimetric cell containing sdAbs at 28 μM. The reference cell was filled with distilled water. In all assays, the titration sequence consisted of a single 0.4 μL injection followed by 19 injections, 2 μL each, with a 150 s time spacing between injections to ensure that the thermal power returned to the baseline before the next injection, and a reference power of 8 μcal/s. The stirring speed was 750 rpm. Control experiments with Ub chains injected in the calorimetric cell filled with buffer were conducted under the same experimental conditions. These control experiments revealed that the heats of dilution were negligible in all cases. The heats per injection normalized per mole of injectant versus the molar ratio [di/tetraUb]/[sdAbs] were fitted by non-linear least-squares regression to a single-site model. Data were analysed with MicroCal PEAQ-ITC (version 1.1.0.1262) analysis software (Malvern Panalytical) and visualised using GraphPad Prism 10.

### Structural modelling

Structural modelling was performed using the AlphaFold3 web interface [43]. M1-linked diUb was modeled as a single protein chain. Predicted Aligned Error (PAE) plots generated from these models were visualised using PAE Viewer [64]. Model confidence was evaluated using standardized metrics including pLDDT, pTM, and ipTM scores, in addition to intermolecular information provided by PAE plots. All structural analysis was performed using PyMol (The PyMOL Molecular Graphics System, Version 3.0 Schrödinger, LLC).

### Protein crystallization, data collection, and processing

Untagged sdAbs 2A6, 2C6, and 2G6 were purified and exchanged into buffer containing 25 mM Tris pH 7.4, 125 mM NaCl, mixed 1:1 with an equivalent concentration of M1-linked diUb, and concentrated to 20 mg/mL. 2C6 was crystallized at 20 °C in a 1 μL sitting-drop containing protein at a 1:1 ratio with 0.2 M calcium chloride dihydrate, 0.1 M HEPES pH 7.5, 28% PEG 400. 2G6 was crystallized at 20 °C in a 1 μL hanging-drop containing protein at a 1:1 ratio with 0.1 M HEPES pH 7.5, and 8% PEG 8000. Both 2C6 and 2G6 crystals were cryoprotected in 25% glycerol prior to vitrification and data collection. 2A6 did not crystallise. Diffraction data were collected on 2C6 crystals at beamline 8.2.2 of the Advanced Light Source Berkeley, CA, while data were collected on 2G6 crystals at beamline 9-2 of the Stanford Synchrotron Radiation Lightsource, Menlo Park, CA. In both cases, the data were integrated using XDS [65] and scaled using either Xtriage [66] (2C6) or AIMLESS [67] (2G6). Structures were determined using the Predict and Build function available in Phenix, which utilises the ColabFold modeling platform for molecular replacement [68–70]. Resulting models underwent iterative rounds of manual model building in Coot [71] and refinement in Phenix [66]. Both 2C6 and 2G6 were found to crystallize on their own, rather than in complex with M1-linked diUb.

### Immunofluorescence and confocal microscopy

HeLa HOIP-KO cells were seeded on cover slips and transiently co-transfected with pcDNA3.1-Htt-Q97-GFP and pcDNA3.1-HA-HOIP variants (WT or C885A) using jetOPTIMUS DNA transfection reagent (Sartorius), according to the manufacturer’s instructions. 48 h after transfection, cells were washed with PBS and then fixed with ice cold methanol (-20 °C) for 30 min. Cells were rehydrated in PBS, permeabilized and blocked using 0.1% Tween 20, 5% BSA, 5% normal goat serum in PBS for 1 h. For sdAb (2A6 and 2E11) staining, the cells were incubated with FLAG-tagged sdAb (20 μg/mL) in the blocking solution overnight at 4oC. Cells were then washed with PBS and incubated anti-FLAG (Cell Signaling) and anti-HA (Cell Signaling) antibody (1:1000) for 3 h at RT. Cells were then washed 3x with PBS and incubated with secondary AlexaFluor-coupled antibodies in PBS for 1 hour at RT. The secondary antibody was removed, and the cover slips washed three times in PBS. All coverslips were incubated in 0.1% (v/v) DAPI in deionised water, washed with deionised water, and then mounted on a glass slide using ProLong Glass Antifade Mountant (Thermo Fisher Scientific). Fluorescence images were acquired using a Leica Stellaris 8 FALCON microscope equipped with an Apo CS2 63x 1.4 NA oil immersion objective. The pinhole was set to 95.5 μm at 1 AU, the samples were scanned at 100 Hz with a pixel size of 0.06 μm, exciting the sample at 405 nm, 499 nm, 568 nm, and 641 nm sequentially. For the objective comparison of fluorescence intensities, confocal images were acquired with the imaging settings kept uniform among replicates and conditions. Following image acquisition, raw data were imported into Imaris 11.0.2 Image analysis software (Andor, Switzerland). Channels were colour adjusted and exported.

### Statistical analysis

Data are presented as mean ± standard deviation (SD). Student’s *t*-test for parametric two-group comparisons was performed using Microsoft Excel. Statistical significance is reported as: ns (not significant, p > 0.05), * (p < 0.05), ** (p < 0.01), *** (p < 0.001).

## Author Contributions

RBD conceived and coordinated the study. JK, S-JB, VB, BKF, BL-M, JNP, and RBD planned experiments. JK, S-JB, VB, BKF, BL-M, and JBR performed experiments and analysed the data. DP, OM-G, MB, GM, MG-H, KFW, and SG contributed reagents or materials. RBD and JK wrote the manuscript. All authors read and approved the manuscript.

## Acknowledgements

We would like to thank Professor J. Preben Morth and Laboratory Technician Suzana Siebenhaar (Technical University of Denmark) for technical assistance. RBD was funded by a Hallas-Møller Emerging Investigator grant from the Novo Nordisk Foundation (NNF19OC005424), and JK was funded by a Walter Benjamin Fellowship from the German Research Association (521996691). Additional support was provided by the NIH National Institute of General Medical Sciences (R35GM142486) to JNP; the LEO Foundation (LF18500) and the Novo Nordisk Foundation (NNF200C0059392) to MG-H; the German Research Foundation (SPP2453, project number 541210481; FOR2848, project number 401510699; RTG2862, project number 492434978) and Germany’s Excellence Strategy (EXC2033 - 390677874 - RESOLV) to KFW; and the Novo Nordisk Foundation (NNF19SA0056783, NNF19SA0057794, and NNF20SA0066621) to SG. Microscopy was supported by the Medical Imaging Center for Light Microscopy (MIC-LM) at Ruhr University Bochum. The funders were not involved in study design, data collection or interpretation, or the writing of this paper.

## Data Availability

A list of key reagents used in this study can be found in Table 1. Further information and requests for reagents should be directed to RBD. Coordinates and structure factors for the 2G6 and 2C6 structures have been deposited in the Protein DataBank under accession codes XXXX and YYYY, respectively, and will be publicly available as of the date of publication.

## AI Disclaimer

Microsoft Copilot was used to improve language and readability of the manuscript. The authors reviewed and edited the content and take full responsibility for the content of the publication.

## Supplementary

**Figure S1. (Related to Figure 1)**

(A) Schematic representation of the structure of the sdAbs in the phage display libraries. CD1 is invariant; CDR2 is variable in sequence but fixed in length; CDR3 is variable in sequence and in length.

(B) Multiple sequence alignment of selected individual sdAb clones identified in this study.

**Figure S2. (Related to Figure 2)**

(A) *In vitro* DUB protection by sdAb 2A6. sdAb 2A6 impairs USP21-mediated hydrolysis of M1-linked tetraUb in a concentration-dependent manner. Reactions were stopped in LSB buffer and run on SDS-PAGE before gels were stained with Coomassie blue.

(B) Overlay of the AlphaFold3 model of 2A6 (grey) with the crystal structures of 2G6 (green) and 2C6 (blue), as well as the HEL4 scaffold (red).

(C) PAE plot from AlphaFold3 of the 2A6 M1 diUb complex shown in Figure 2D.

(D) AlphaFold3 model of the 2A6 M1 diUb complex colored by pLDDT confidence scores.

(E) Overlay of the five top scoring AlphaFold3 models of the 2A6 M1 diUb complex.

(F) List of positions in sdAb 2A6 subjected to saturation mutagenesis as well as the identified permitted variants. Positions are coloured based on the conservation of permitted variants: green, non-conservative substitutions tolerated; yellow, conservative substitutions permitted; red, no substitutions permitted.

(G) ELISA of selected 2A6 variants from saturation mutagenesis expressed in the bacterial periplasm, showing binding to K48-, K63-, and M1-linked diUb. Bars represent mean. Dashed lines indicate the signal obtained by two independent samples of parental 2A6.

**Figure S3. (Related to Figure 3)**

(A) Ponceau S-stained membranes from the experiment in Figure 3D show the IP or pulldown levels of sdAb 2A6, M1-Trap, and M1-SUB.

**Figure S4. (Related to Figure 4)**

(A) Immunoblots of IP with His-tagged sdAb 2E11 from control (OTULIN^+/-^) and OTULIN-KO MEFs in IP buffer without NaDC and SDS detergents.

**Table S1:** X-ray crystallography data collection and refinement statistics.

|  | 2G6 | 2C6 |
| --- | --- | --- |
| **Data collection** |  |  |
| Wavelength | 0.979458 | 1.000 |
| Space group | *I* 1 2 1 | *P* 6<sub>3</sub> 2 2 |
| **Cell dimensions** |  |  |
| *a*, *b*, *c* (Å) | 30.07, 70.90, 67.12 | 64.71, 64.71, 102.68 |
| $\alpha$, $\beta$, $\gamma$ (°) | 90, 102.36, 90 | 90, 90, 120 |
| Resolution (Å) | 48.13-1.40 (1.42-1.40) | 49.19-2.65 (2.78-2.65) |
| *R*<sub>merge</sub> | 0.037 (0.407) | 0.319 (1.886) |
| *I*/σ*I* | 21.6 (3.1) | 7.9 (1.5) |
| Completeness (%) | 93.6 (79.2) | 100.0 (100.0) |
| Redundancy | 6.8 (5.5) | 9.3 (10.0) |
| **Phasing** |  |  |
| Method | MR | MR |
| MR Search Models | AF3 model | AF3 model |
| **Refinement** |  |  |
| Resolution (Å) | 48.13-1.40 | 49.19-2.65 |
| No. unique reflections / test set | 25291 / 1277 | 4063 / 222 |
| *R*<sub>work</sub>/*R*<sub>free</sub> | 0.181 / 0.217 | 0.302 / 0.360 |
| **No. atoms** |  |  |
| Protein | 929 | 881 |
| Ligand/ion | 12 | 0 |
| Water | 146 | 0 |
| **B-factors** |  |  |
| Protein | 20.76 | 44.68 |
| Ligand/ion | 45.15 | - |
| Water | 33.30 | - |
| **R.m.s. deviations** |  |  |
| Bond lengths (Å) | 0.013 | 0.005 |
| Bond angles (°) | 1.33 | 0.87 |
| **Ramachandran statistics** |  |  |
| Favored / allowed / outliers | 99.2 / 0.8 / 0 | 90.3 / 8.8 / 0.9 |

Values in parentheses are for highest resolution shell. AF3, AlphaFold3

