## Supplemental Figures for "Development and Characterisation of a Versatile Single-Domain Antibody Specific for M1-linked Ubiquitin Chains"

### Figure S1

## A

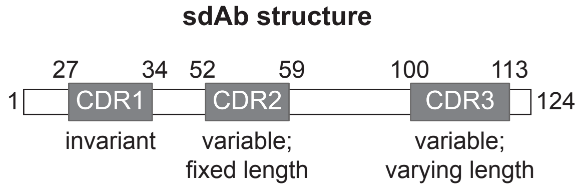


## B

**Sequence alignment of selected clones:**

```
             2                        27     34                     52     59                                      100          113        124
Consensus    EVQLLESGGGLVQPGGSLRLSCAASGFRISDEDMGWFRQAPGKEREWVSSISSRFGNTYYADSVKGRFTISRDNSKNTLYLQMNSLRAEDTAVYYCARGYRDVDFLDYSFDVWGQGTQVTVSS
1H8          EVQLLESGGGLVQPGGSLRLSCAASGFRISDEDMGWFRQAPGKEREWVSSISSIFGNKYYADSVKGRFTISRDNSKNTLYLQMNSLRAEDTAVYYCARERGTPY---YGFDVWGQGTQVTVSS
1A3          EVQLLESGGGLVQPGGSLRLSCAASGFRISDEDMGWFRQAPGKEREWVSSISPRFGNTYYADSVKGRFTISRDNSKNTLYLQMNSLRAEDTAVYYCARGVR---FFDYFYDSWGQGTQVTVSS
1C5          EVQLLESGGGLVQPGGSLRLSCAASGFRISDEDMGWFRQAPGKEREWVSSISPRFGNTYYADSVKGRFTISRDNSKNTLYLQMNSLRAEDTAVYYCARGVR---FFDYFYDSWGQGTQVTVSS
1G6          EVQLLESGGGLVQPGGSLRLSCAASGFRISDEDMGWFRQAPGKEREWVSSISYSFGNTYYADSVKGRFTISRDNSKNTLYLQMNSLRAEDTAVYYCARERSYPTNGTYAFDVWGQGTQVTVSS
1A8          EVQLLESGGGLVQPGGSLRLSCAASGFRISDEDMGWFRQAPGKEREWVSSISPIYGNTYYADSVKGRFTISRDNSKNTLYLQMNSLRAEDTAVYYCARGYR---LYSGTYDVWGQGTQVTVSS
1E9          EVQLLESGGGLVQPGGSLRLSCAASGFRISDEDMGWFRQAPGKEREWVSSISSIFGNKYYADSVKGRFTISRDNSKNTLYLQMNSLRAEDTAVYYCARRVYRVG---SYYDYWGQGTQVTVSS
1A1          EVQLLESGGGLVQPGGSLRLSCAASGFRISDEDMGWFRQAPGKEREWVSSITARYRNTYYADSVKGRFTISRDNSKNTLYLQMNSLRAEDTAVYYCARAYGRTD---YEYDVWGQGTQVTVSS
1E11         EVQLLESGGGLVQPGGSLRLSCAASGFRISDEDMGWFRQAPGKEREWVSSIKWIFGNTYYADSVKGRFTISRDNSKNTLYLQMNSLRAEDTAVYYCARGYGGGGGYGYEFDVWGQGTQVTVSS
1H11         EVQLLESGGGLVQPGGSLRLSCAASGFRISDEDMGWFRQAPGKEREWVSSIKWIFGNTYYADSVKGRFTISRDNSKNTLYLQMNSLRAEDTAVYYCARGYGGGGGYGYEFDVWGQGTQVTVSS
2B1          EVQLLESGGGLVQPGGSLRLSCAASGFRISDEDMGWFRQAPGKEREWVSSISWSYGNTYYADSVKGRFTISRDNSKNTLYLQMNSLRAEDTAVYYCARGYDGSI---YSFDYWGQGTQVTVSS
2G1          EVQLLESGGGLVQPGGSLRLSCAASGFRISDEDMGWFRQAPGKEREWVSSISSRYDNTYYADSVKGRFTISRDNSKNTLYLQMNSLRAEDTAVYYCARAYGRTD---YEYDVWGQGTQVTVSS
2F2          EVQLLESGGGLVQPGGSLRLSCAASGFRISDEDMGWFRQAPGKEREWVSSISGRFSNKYYADSVKGRFTISRDNSKNTLYLQMNSLRAEDTAVYYCARRYGDGDLDDYTFDVWGQGTQVTVSS
2B3          EVQLLESGGGLVQPGGSLRLSCAASGFRISDEDMGWFRQAPGKEREWVSSISSIFGNKYYADSVKGRFTISRDNSKNTLYLQMNSLRAEDTAVYYCARGVRDVSFLDYSFDSWGQGTQVTVSS
2E4          EVQLLESGGGLVQPGGSLRLSCAASGFRISDEDMGWFRQATGKEREWVSSISSIYSNAYYADSVKGRFTISRDNSKNTLYLQMNSLRAEDTAVYYCARGVRDVSFLDYSFDSWGQGTQVTVSS
2B5          EVQLLESGGGLVQPGGSLRLSCAASGFRISDEDMGWFRQAPGKEREWVSSITASTGNIYYADSVKGRFTISRDNSKNTLYLQMNSLRAEDTAVYYCARGVRDVSFLDYSFDSWGQGTQVTVSS
2A6          EVQLLESGGGLVQPGGSLRLSCAASGFRISDEDMGWFRQAPGKEREWVSSISGRFSNKYYADSVKGRFTISRDNSKNTLYLQMNSLRAEDTAVYYCARGLPGPD---YSFDVWGQGTQVTVSS
2C6          EVQLLESGGGLVQPGGSLRLSCAASGFRISDEDMGWFRQAPGKEREWVSSISGRFSNKYYADSVKGRFTISRDNSKNTLYLQMNSLRAEDTAVYYCARRYGDGDLDDYTFDVWGQGTQVTVSS
2G6          EVQLLESGGGLVQPGGSLRLSCAASGFRISDEDMGWFRQAPGKEREWVSSISSIFGNKYYADSVKGRFTISRDNSKNTLYLQMNSLRAEDTAVYYCARGVRDVSFLDYSFDSWGQGTQVTVSS
2A7          EVQLLESGGGLVQPGGSLRLSCAASGFRISDEDMGWFRQAPGKEREWVSSIYPTFGNTYYADSVKGRFTISRDNSKNTLYLQMNSLRAEDTAVYYCARRYGDGDLDDYTFDVWGQGTQVTVSS
2A10         EVQLLESGGGLVQPGGSLRLSCAASGFRISDEDMGWFRQAPGKEREWVSSISGRFSNKYYADSVKGRFTISRDNSKNTLYLQMNSLRAEDTAVYYCARRYGDGDLDDYTFDVWGQGTQVTVSS
2F3          EVQLLESGGGLVQPGGSLRLSCAASGFRISDEDMGWFRQAPGKEREWVSSISSRYDNTYYADSVKGRFTISRDNSKNTLYLQMNSLRAEDTAVYYCARAYGRTD---YEYDVWGQGTQVTVSS
2C12         EVQLLESGGGLVQPGGSLRLSCAASGFRISDEDMGWFRQAPGKEREWVSSISSRYDNTYYADSVKGRFTISRDNSKNTLYLQMNSLRAEDTAVYYCARAYGRTD---YEYDVWGQGTQVTVSS
2E12         EVQLLESGGGLVQPGGSLRLSCAASGFRISDEDMGWFRQAPGKEREWVSSISSRYDNTYYADSVKGRFTISRDNSKNTLYLQMNSLRAEDTAVYYCARERRPSYDGGYYFDVWGQGTQVTVSS
2G12         EVQLLESGGGLVQPGGSLRLSCAASGFRISDEDMGWFRQAPGKEREWVSSISSRYDNTYYADSVKGRFTISRDNSKNTLYLQMNSLRAEDTAVYYCARAYGRTD---YEYDVWGQGTQVTVSS
R3/P2        EVQLLESGGGLVQPGGSLRLSCAASGFRISDEDMGWFRQAPGKEREWVSSISSIFGNKYYADSVKGRFTISRDNSKNTLYLQMNSLRAEDTAVYYCARGFERTD---YEYDVWGQGTQVTVSS
RPA4         EVQLLESGGGLVQPGGSLRLSCAASGFRISDEDMGWFRQAPGKEREWVSSVSSRDSTSYYADSVKGRFTISRDNSKNTLYLQMNSLRAEDTAVYYCARGWNGSGFRPYEFDYWGQGTQVTVSS
RPA8         EVQLLESGGGLVQPGGSLRLSCAASGFRISDEDMGWFRQAPGKEREWVSSINGDFGNKYYADSVKGRFTISRDNSKNTLYLQMNSLRAEDTAVYYCARGFRGDVLYSYSFDVWGQGTQVTVSS
RPA15        EVQLLESGGGLVQPGGSLRLSCAASGFRISDEDMGWFRQAPGKEREWVSSISASFSNPYYADSVKGRFTISRDNSKNTLYLQMNSLRAEDTAVYYCARGLPGPD---YSFDVWGQGTQVTVSS
PBS1         EVQLLESGGGLVQPGGSLRLSCAASGFRISDEDMGWFRQAPGKEREWVSSIYRTSGNTYYADSVKGRFTISRDNSKNTLYLQMNSLRAEDTAVYYCARYYRPRP---RYFDVWGQGTQVTVSS
PBS3         EVQLLESGGGLVQPGGSLRLSCAASGFRISDEDMGWFRQAPGKEREWVSSINWQYGSIYYADSVKGRFTISRDNSKNTLYLQMNSLRAEDTAVYYCARALSYDS---LSYDVWGQGTQVTVSS
PBS5         EVQLLESGGGLVQPGGSLRLSCAASGFRISDEDMGWFRQAPGKEREWVSSISGRFSNKYYADSVKGRFTISRDNSKNTLYLQMNSLRAEDTAVYYCARALRGDP---RYFDVWGQGTQVTVSS
PBS6         EVQLLESGGGLVQPGGSLRLSCAASGFRISDEDMGWFRQAPGKEREWVSSISRGFDTTYYADSVKGRFTISRDNSKNTLYLQMNSLRAEDTAVYYCARGYRTRASYIGYFDIWGQGTQVTVSS
PBS9         EVQLLESGGGLVQPGGSLRLSCAASGFRISDEDMGWFRQAPGKEREWVSSISGRFSNKYYADSVKGRFTISRDNSKNTLYLQMNSLRAEDTAVYYCARRWSYPGAGSRAFDVWGQGTQVTVSS
PBS16        EVQLLESGGGLVQPGGSLRLSCAASGFRISDEDMGWFRQAPGKEREWVSSISSIFGNKYYADSVKGRFTISRDNSKNTLYLQMNSLRAEDTAVYYCARSLPVSI---YSYDVWGQGTQVTVSS
2A6.1        EVQLLESGGGLVQPGGSLRLSCAASGFRISDEDMGWFRQAPGKEREWVSSISSIFGNKYYADSVKGRFTISRDNSKNTLYLQMNSLRAEDTAVYYCARGLPGPD---YSFDVWGQGTQVTVSS
2G6.1        EVQLLESGGGLVQPGGSLRLSCAASGFRISDEDMGWFRQAPGKEREWVSSISGRFSNKYYADSVKGRFTISRDNSKNTLYLQMNSLRAEDTAVYYCARGVRDVSFLDYSFDSWGQGTQVTVSS
2E11         EVQLLESGGGLVQPGGSLRLSCAASGFRISDEDMGWFRQAPGKEREWVSSISSSFGNTYYADSVKGRFTISRDNSKNTLYLQMNSLRAEDTAVYYCARELRRGSPIEYWYDVWGQGTQVTVSS
1E3 (HEL4)   EVQLLESGGGLVQPGGSLRLSCAASGFTFSNTDMGWFRQAPGKEREWVSSITPSSGQTYYADSVKGRFTISRDNSKNTLYLQMNSLRAEDTAVYYCARTWLL----RWVMDLWGQGTQVTVSS
2E4.1        EVQLLESGGGLVQPGGSLRLSCAASGFRISDEDMGWFRQATGKEREWVSSISSIYSQAYYADSVKGRFTISRDNSKNTLYLQMNSLRAEDTAVYYCARGVRDVSFLDYSFDSWGQGTQVTVS-
2G9          EVQLLESGGGLVQPGGSLRLSCAASGFRISDEDMGWFRQAPGKEREWVSSISSIYSNAYYADSVKGRFTISRDNSKNTLYLQMNSLRAEDTAVYYCARGVRDVSFLDYSFDSWGQGTQVTVSS
```

### Figure S2

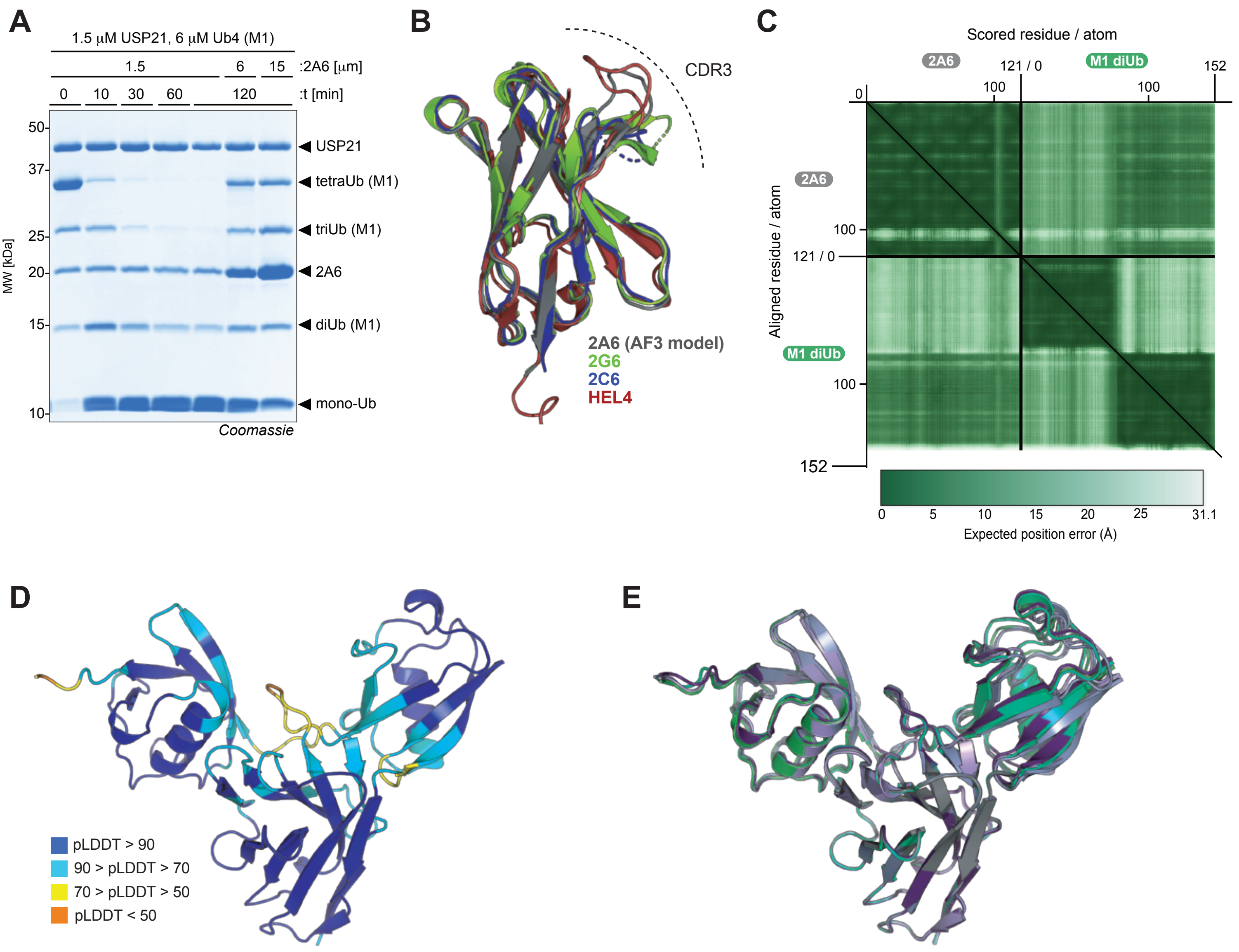


F

##### Saturation mutagenesis

| Position | Permitted variants | Classification |
|---|---|---|
| F38 | Y | Conservative substitution tolerated |
| Q40 | W | Non-conservative substitution tolerated |
| E45 | R | Non-conservative substitution tolerated |
| R46 | Y | Non-conservative substitution tolerated |
| E47 | T, K, S, M | Non-conservative substitution tolerated |
| W48 | | No substitutions tolerated |
| S51 | K | Non-conservative substitution tolerated |
| R55 | K | Conservative substitution tolerated |
| F56 | Y | Conservative substitution tolerated |
| N58 | H | Conservative substitution tolerated |
| Y60 | | No substitutions tolerated |
| A62 | R | Non-conservative substitution tolerated |
| D63 | E, S, H | Non-conservative substitution tolerated |
| K66 | | No substitutions tolerated |
| L101 | V, F, Y, C | Non-conservative substitution tolerated |
| D105 | | No substitutions tolerated |
| Y106 | | No substitutions tolerated |
| S107 | | No substitutions tolerated |
| F108 | | No substitutions tolerated |
| D109 | | No substitutions tolerated |
| V110 | L | Conservative substitution tolerated |
| W111 | T, R, V, I | Non-conservative substitution tolerated |

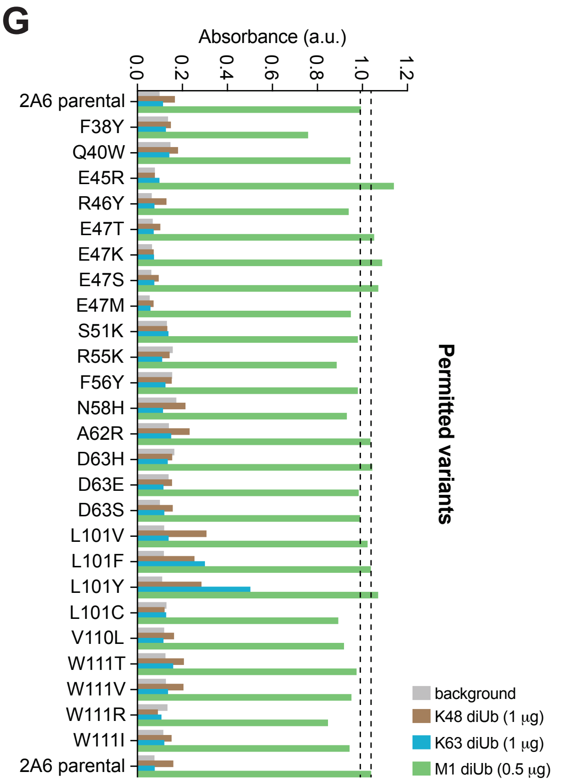

### Figure S3

## A

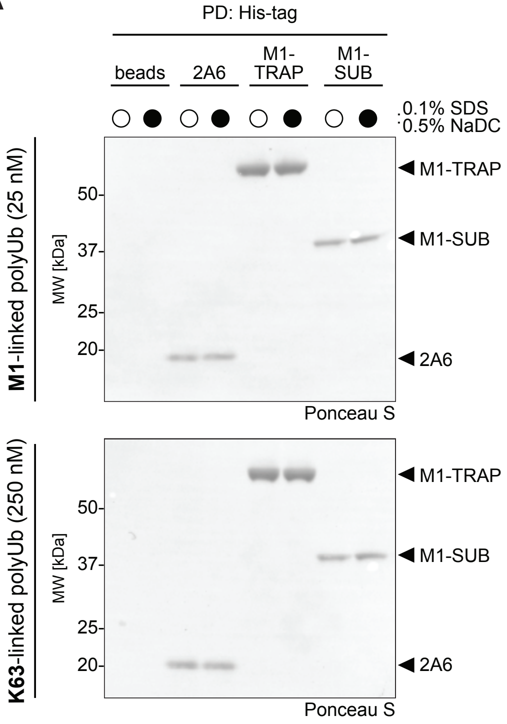

PD: His-tag
beads
2A6
M1-TRAP
M1-SUB
0.1% SDS
0.5% NaDC
M1-linked polyUb (25 nM)
MW [kDa]
50-
37-
25-
20-
M1-TRAP
M1-SUB
2A6
Ponceau S
K63-linked polyUb (250 nM)
MW [kDa]
50-
37-
25-
20-
M1-TRAP
M1-SUB
2A6
Ponceau S

### Figure S4

**A**

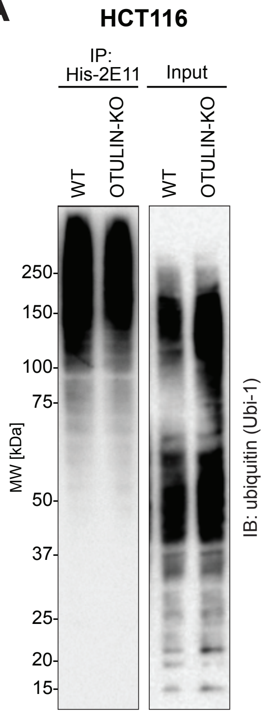
HCT116
IP: His-2E11
Input
WT
OTULIN-KO
WT
OTULIN-KO
MW [kDa]
250
150
100
75
50
37
25
20
15
IB: ubiquitin (Ubi-1)
